# Quantifying signal persistence in the T cell signaling network using an optically controllable antigen receptor

**DOI:** 10.1101/2020.10.30.362194

**Authors:** Michael J Harris, Muna Fuyal, John R James

## Abstract

T cells discriminate between healthy and infected cells with remarkable sensitivity when mounting an immune response. It has been hypothesized that this efficient detection requires combining signals from discrete antigen-presenting cell interactions into a more potent response, requiring T cells to maintain a ‘memory’ of previous encounters. To quantify the magnitude of this phenomenon, we have developed an antigen receptor that is both optically and chemically tunable, providing control over the initiation, duration, and intensity of intracellular T-cell signaling within physiological cell conjugates. We observe very limited persistence within the T cell intracellular network on disruption of receptor input, with signals dissipating entirely in ~15 minutes, and directly confirm that sustained proximal receptor signaling is required to maintain active gene transcription. Our data suggests that T cells are largely incapable of integrating discrete antigen receptor signals but instead simply accumulate the output of gene expression. By engineering optical control in a clinically relevant chimeric antigen receptor, we show that this limited signal persistence can be exploited to increase the activation of primary T cells by ~3-fold by using pulsatile stimulation. Our results are likely to apply more generally to the signaling dynamics of other cellular networks.

## INTRODUCTION

The ability of cells to convert extracellular stimuli into information that can guide future decisions is an essential requirement for organisms to survive within their environment (Jordan et al., 2000). External inputs are sensed by receptors that reside predominantly at the cell surface in mammalian cells, whereas their functional output is normally manifested within the cell nucleus as changes in gene expression. This output can drive long term alterations in cell function or activity by bringing about a new state that persists when the originating input has been removed and constitutes a memory of previous signaling (Burrill and Silver, 2010). The transduction of signals from the surface receptors to the transcriptome is mediated by an intracellular network of signaling proteins and lipids that transmit and amplify the cellular inputs. While it is well established that signaling networks have significant information processing capabilities (Atay and Skotheim, 2017; De et al., 2020; Martin et al., 2020), less is known about signal persistence within such networks when the initiating stimulus is removed; a quantitative understanding of this response requires elucidating the timescales of how intermediary signals decay within the network. Examples of this may include the phosphorylation of a protein or the active state of a promoter. Individual steps in an intracellular signaling network are invariably counteracted by opposing biochemical reactions; how efficient these reverse steps are will define the level of signal persistence within the network. Any potential for signals to be retained on cessation of receptor input would constitute a short-term memory, which could be observed directly when distinct inputs are separated over time providing a mechanism to combine discrete events into a more potent response compared to the individual pulses in isolation.

T cell activation is a physiological situation where integrating multiple signaling events could drive a more robust output response. T cells are an essential cell-type of our adaptive immune system that keep us healthy by continually scanning the surface of other cells looking for signs of infection. The T-cell antigen receptor (TCR) is a multi-protein complex expressed within the plasma membrane that is the primary mediator of this immune surveillance. The TCR can recognize pathogen-derived peptides in the MHC protein complex (pMHC) expressed on almost all cells and the information encoded in this interaction is relayed to the T cell interior, initiating a signaling cascade that ultimately leads to changes in T cell activation and cell-fate differentiation (Brownlie and Zamoyska, 2013; Smith-Garvin et al., 2009). Importantly, T cells are known to be exceptionally sensitive, with the ability to show a detectable response to cells presenting just a few cognate pMHC ligands (Anikeeva et al., 2012; Huang et al., 2013).

T cells continuously interact with other cells as they traverse between the blood and lymph circulation systems, primarily engaging with antigen-presenting cells (APC) in lymph nodes and forming mobile synapses, termed kinapses (Dustin, 2008). Previous work *in vitro* has shown that T cells can be activated over multiple APC interactions even though the individual interactions are likely to be transient (Gunzer et al., 2000; Underhill et al., 1999), results that have been corroborated from intravital imaging in mice (Celli et al., 2005; Le Borgne et al., 2016; Marangoni et al., 2013). However, it is also been found that sustained proximal signaling is required to drive effective T cell activation (Au-Yeung et al., 2014; Smith-Garvin et al., 2009; Valitutti et al., 1995), which would require T cells to combine sub-optimal signaling events initiated during these transient encounters, known as ‘Phase-One’ (Mempel et al., 2004), into more robust activation. Periodic signal disruption has shown that T cells are capable of combining temporally separated stimuli (Clark et al., 2011; Faroudi et al., 2003; Munitic et al., 2005) but may affect the differentiation pathway of the activated T cells when serial transient interactions are formed (Scholer et al., 2008).

To investigate this discrepancy between the requirement for sustained signaling and interactions with multiple APC, we sought to quantify the rate at which receptor signaling dissipates within T cells to define the timescales over which the intracellular network has the capacity to integrate the signals from discrete receptor inputs. Exciting new methods to do this have come from the development of optogenetic toolkits that provide the means to control signaling in a non-invasive and reversible manner (Bugaj et al., 2017; Toettcher et al., 2013). Recent studies have used these approaches to understand the dynamics of signaling within the ERK pathway and how this information is encoded at the transcriptional level (Wilson et al., 2017). There have also been two studies that have used optically controllable ligands to investigate whether the TCR can readout the lifetime of the TCR/pMHC interaction. However, these results were achieved with either soluble tetrameric (Yousefi et al., 2019) or bilayer-bound (Tischer and Weiner, 2019) ligands, which may not be capable of driving persistent downstream signaling as they remove the physiological context of the T cell-APC interface and other costimulatory signals. Another recent study has demonstrated that the initiation of calcium fluxing in T cells can be controlled *in vivo* using an optogenetic actuator (Bohineust et al., 2020).

We have developed a TCR-based receptor where signaling can be synchronously initiated within T cell-APC conjugates combined with optical control over the signaling dynamics. We used this tool to quantify the persistence of signals at representative intracellular nodes within the TCR signal transduction network when receptor input is acutely disrupted. Through this approach, we have found that intracellular signals dissipate completely within ~15 minutes, with mRNA levels providing the most persistent intermediary state with a half-life of ~25 minutes. It is therefore unlikely that T cells directly integrate TCR signals between multiple APC interaction, but rather accumulate the transcriptional output of gene expression from each stimulation. We also show that this limited signal persistence can be exploited to drive more efficient signaling in primary T cells by transplanting optical control to a clinically relevant chimeric antigen receptor.

## RESULTS

### Construction of an optically modulated chimeric antigen receptor in T cells

Our first goal was to design a synthetic antigen receptor in T cells that would provide a stimulus to the intracellular T cell signaling network that was under light-mediated control. Furthermore, this optically modulated receptor needed to function within the context of the interface between a T cell and APC. This requirement is important as it allows other signaling pathways to function unperturbed and most closely mimics the native signaling environment. The most efficient way to achieve this was by engineering in the ability to physically uncouple extracellular ligand binding from signal transduction across the plasma membrane (Figure 1A). This was accomplished using the LOVTRAP system (Wang et al., 2016), where illumination of a LOV2 domain with blue light (400-500nm) causes the reversible dissociation of an engineered Zdk domain, which only binds the dark state of LOV2. To create our optically controlled chimeric antigen receptor, which we term an ‘OptoCAR’, we fused the light sensitive LOV2 domain to the intracellular terminus of a synthetic receptor that we have previously shown replicates the function of the native TCR complex (James, 2018; James and Vale, 2012). The ITAM signaling motifs from the TCR complex (CD3ζ) were fused to the Zdk domain and this intracellular part of the OptoCAR was constrained to the plasma membrane through myristoylation of its N-terminus (Figure 1B). This configuration meant that on light-mediated dissociation of Zdk from the transmembrane component of the receptor, the signaling moiety would diffuse away laterally from the receptor-ligand complex and be rapidly dephosphorylated by CD45 phosphatase, which most closely replicates the response to dissociation of the TCR from its pMHC ligand (Figure 1B).

**Figure 1.**
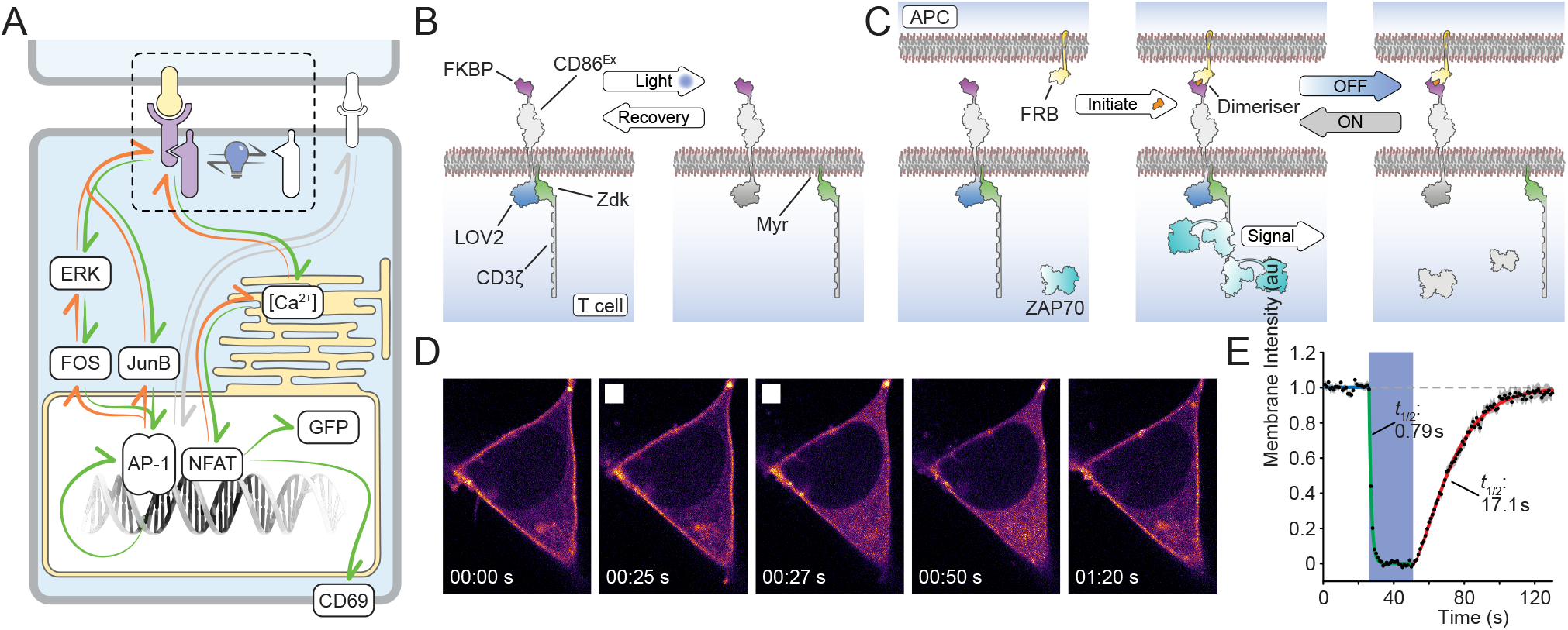
Construction of an optically modulated chimeric antigen receptor (OptoCAR) in T cells. (A) Schematic showing the OptoCAR replacing the antigen receptor (dotted region) as the primary driver of intracellular T cell signaling. Illumination of the cells leads to the dissociation of the intracellular part of the OptoCAR and the loss of signal transduction. The network nodes investigated in this study are explicitly depicted within the boxes. (B) A more detailed schematic of the bipartite OptoCAR from (A) with component parts labelled. Light-mediated disruption causes dissociation of intracellular signaling tail (ζ-chain) from the extracellular binding domain. (C) OptoCAR-T cell engagement with an antigen-presenting cell (APC) does not initiate signaling in the absence of the dimerizer small-molecule. OptoCAR signaling from within these cell conjugates is modulated by transitioning from the dark (signaling competent) state to the ‘Off’ state by illuminating cells with blue light. The intracellular domain will be rapidly dephosphorylated when it diffuses away from the bound OptoCAR. (D) Stills from Movie S1 showing the membrane-bound form of the OptoCAR expressed in HEK293T cells being reversibly dissociated from the extracellular domain by single-cell illumination (white box). The intracellular part of the OptoCAR used in this experiment was not myristoylated so it localized to the cytoplasm on dissociation, to aid visualization of OptoCAR dynamics. (E) Quantification of the light-mediated dissociation of OptoCAR in HEK293T cells by measuring the ratio of membrane-bound intracellular part of OptoCAR over time. Individual cells were illuminated after 25 seconds of imaging for 25 seconds to calculate dissociation rate (green) before returning to dark (signaling competent) state to calculate re-association rate (red line). Bounding area around datapoints shows mean ± SEM (n=12 cells).

The ectodomain of the OptoCAR described above provides an orthogonal T cell input that can be initiated on addition of a small molecule dimerizer drug (Figure 1C). The power of this approach is that it allows the process of cell-cell engagement and interface formation to be temporally separated from the initiation of receptor signaling. Dimerizer addition can then initiate signaling within the context of T cell-APC conjugates in a synchronized manner, which is essential to follow signaling dynamics at high temporal resolution. The integration of optical and chemical inputs within the OptoCAR presented a unique way to control both the intensity and duration of proximal signaling at the T cell conjugate interface and follow the associated downstream response (Figure 1C). Importantly, optical control does not directly perturb any part of the binding interface between the two cells and so engagement of any other co-stimulatory receptors, such as CD28, would remain unaffected.

We first wanted to directly test the response of the OptoCAR to light control. We expressed the receptor in HEK-293T cells, which are non-immune cells and do not express proteins that normally interact with the TCR. In its normal bipartite architecture, the dissociation of the OptoCAR on light stimulation would simply alter the diffusion rate of the two components, which is difficult to quantify. We therefore modified the OptoCAR so that the intracellular fragment would dissociate and diffuse freely in the cytoplasm. This then allowed us to visualize the light-mediated dissociation of the OptoCAR by labelling the intracellular fragment with a red fluorescent protein (mScarlet) and measuring its cellular localization by confocal microscopy.

Stimulation with blue light caused the rapid dissociation of the bipartite receptor (Figures 1D, 1E and Movie S1) with an off-rate of 0.88 s^−1^, implying near-complete (95%) dissociation within ~3.5 seconds. Turning off the light causes the thermally induced reversion to the ‘dark’ (signaling-competent) state of the LOV2 domain and hence reformation of the signal-competent OptoCAR (Figures 1D, 1E and Movie S1). This process was slower with an on-rate of 0.04 s^−1^ and near-complete complex association within ~75 seconds, which agrees very well with previously reported values for the wildtype LOVTRAP system (Wang et al., 2016).

### Activation-induced intracellular Ca^2+^ flux is rapidly disrupted on signal cessation

Having shown that the OptoCAR itself was both light responsive and could be reversibly modulated, we next wanted to test whether the OptoCAR functioned as anticipated in T cells to influence the proximal receptor signaling pathway. We used viral transduction to stably express the OptoCAR in the Jurkat T cell line (OptoCAR-T cells) and confirmed that the synthetic reporter was readily detectable at the cell surface (Figures S1A-C).

One of the earliest direct readouts of T cell activation is Ca^2+^ fluxing, the rapid increased concentration of Ca^2+^ ions in the cytoplasm from ER-derived stores. T cells expressing the OptoCAR were first loaded with indo-1, a ratiometric Ca^2+^ sensor, before being conjugated with a B cell line expressing the counterpart ligand (Raji-FRB^Ex^). Pertinently, the initial interaction between the two cell types was driven solely by adhesion proteins in the absence of the dimerizer, so that proximal signaling remained uninitiated. Flow cytometry was then used to gate on only these cell conjugates prior to dimerizer addition (Figures S1D and S1E) and follow changes in intracellular Ca^2+^ concentration over time.

We first confirmed that no cellular response was detectable when a vehicle control rather than the dimerizer was used to initiate OptoCAR-mediated signaling (Figure 2A). We then performed the assay in the ‘dark’ state (without illumination) to assess whether OptoCAR engagement could initiate proximal T cell signaling. The addition of dimerizer to the conjugated T cells did indeed cause a robust increase in intracellular concentration of Ca^2+^ ions, with the flux saturating within ~2 minutes (Figure 2B).

**Figure 2.**
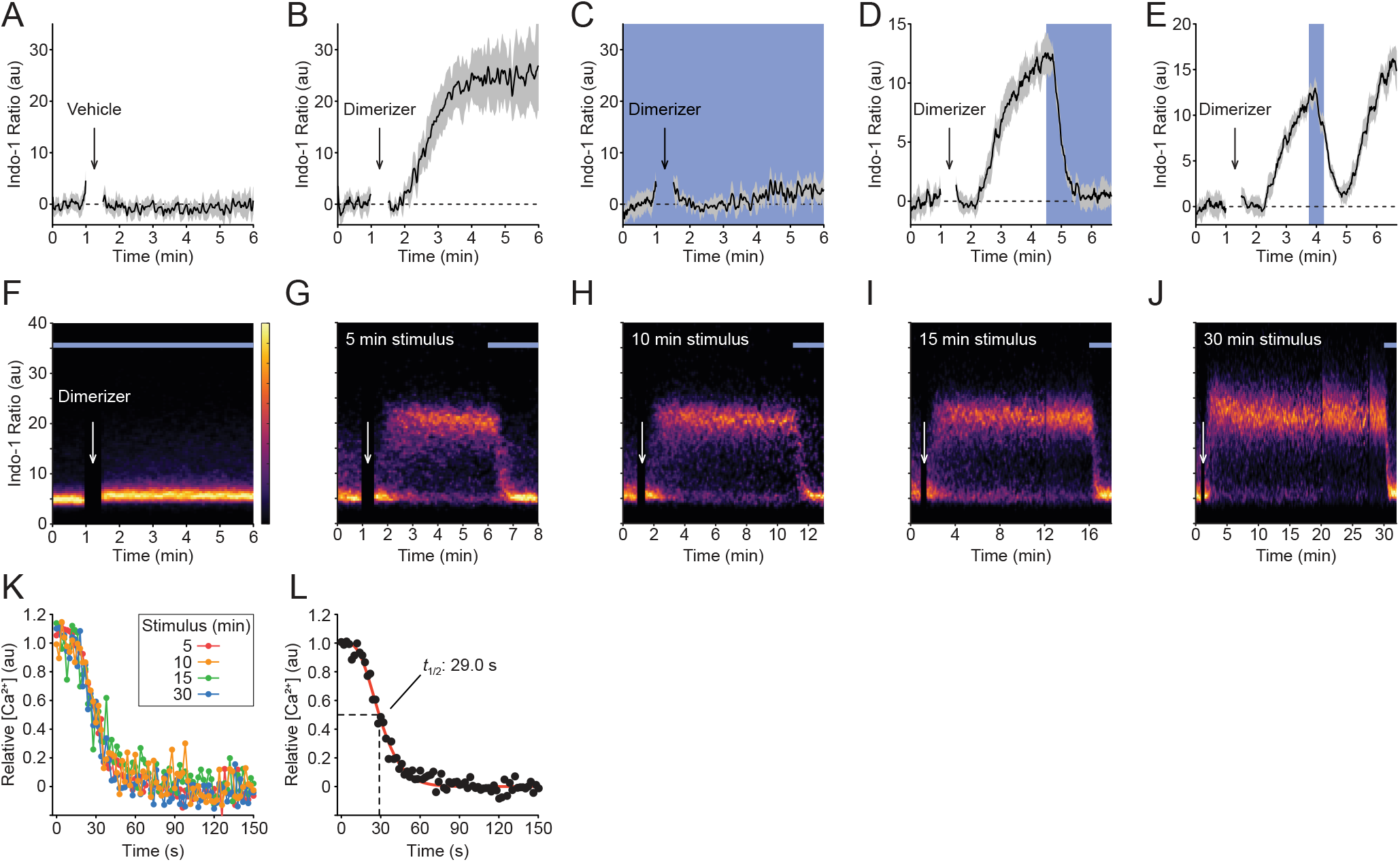
Activation-induced intracellular Ca^2+^ flux is rapidly disrupted on signal cessation. (A) OptoCAR-T cells were loaded with a ratiometric Ca^2+^indicator (Indo-1) before conjugation with ligand-presenting cells. Conjugated cells were gated on by flow cytometry and the Indo-1 ratio, as a readout of intracellular [Ca^2+^] was measured over time. Vehicle addition after 60 seconds caused no detectable change in this ratio. (B) An equivalent experiment was setup as in (A) but now 2 μM dimerizer was added after 60 seconds. Conjugated cells were maintained in the dark to maintain OptoCAR signaling. (C) An equivalent experiment was setup as in (B) but the conjugated cells were continuously illuminated throughout dimerizer addition to disrupt OptoCAR signaling. (D) An equivalent experiment was setup as in (B) but now conjugated cells were illuminated 3.5 minutes after dimerizer addition. (E) An equivalent experiment was setup as in (B) but now cells were illuminated 165 seconds after dimerizer addition, for 30 seconds. (F-J) Representative density plots of Ca^2+^ flux from conjugated OptoCAR-T cells over time at 37°C. Conjugated cells were stimulated with dimerizer addition for 0 (F), 5 (G), 10 (H), 15 (I) or 30 (J) minutes prior to disrupting signaling by illuminating cells. (K) The decrease in intracellular [Ca^2+^] on illuminating conjugated cells after different intervals during activation was plotted as the mean Indo-1 ratio (n=3). Inset legend delineates the datasets. (L) A single plot combining all datasets from (K) was fit by the survival function of a gamma distribution, with indicated half-life calculated from the time taken for the relative [Ca^2+^] to decrease to 50% of maximum. See also Figure S1.

To provide optical control over receptor signaling whilst simultaneously measuring the calcium flux, we custom-built a heated LED illuminator that could be installed directly onto the flow cytometer (Figure S1F). Repeating the assay above under constant blue light illumination completely inhibited Ca^2+^ fluxing to an equivalent extent as the vehicle control (Figure 2C), showing that the OptoCAR was indeed light responsive in T cells. To rule out that this observation was due to any phototoxic effects of light stimulation we repeated the assay with a LOV2 domain mutant (C450G) that is unresponsive to light (Wang et al., 2016). In this case, the observed Ca^2+^ flux was found to be maintained even under constant illumination (Figure S1G). Conversely, using a LOV2 mutant (I539E) that remains open even in the dark (Wang et al., 2016), we could not observe any fluxing (Figure S1H), confirming that the OptoCAR was not simply creating a signaling competent region within the plasma membrane for the endogenous TCR (or other receptors) at the cell surface to signal.

Having convinced ourselves that the OptoCAR was responsive to light input, we wanted to use our new tool to investigate the kinetics and reversibility of how proximal signaling drives Ca^2+^ fluxing. We repeated the experiment above with the wildtype OptoCAR, but this time illuminated the cells once the intracellular concentration of Ca^2+^ ion had begun to increase upon dimerizer addition. Illumination of the cells undergoing proximal signaling caused the rapid termination of the Ca^2+^ flux, which reverted to baseline within approximately 30 seconds (Figure 2D). This light-mediated disruption of proximal signaling was readily reversible once illumination ceased and the OptoCAR reverted to the signaling-competent state (Figure 2E).

The extremely rapid decrease in intracellular Ca^2+^ concentration on light-mediated cessation of receptor signaling was unexpected given that illumination of cells only disrupts the functional state of the receptor itself. We therefore questioned whether this result was because we had disrupted signaling so soon after its initiation and had not allowed proximal signaling to become self-sustaining, if it had this capacity. To address this point, we performed the equivalent assay but illuminated the conjugates to disrupt proximal signaling at different timepoints after its initiation. Constant illumination caused no detectable Ca^2+^ flux in the raw cytometry plots as expected (Figure 2F) but light-mediated cessation of receptor activation caused a rapid decrease in Ca^2+^ fluxing even after 30 minutes activation (Figures 2G – 2J). We quantified the dynamics of Ca^2+^ flux disruption on illumination and found that all stimulation periods showed essentially equivalent kinetics of signal decay (Figure 2K). This implied that this part of the proximal signaling network remains rapidly reversible and is entirely contingent on continuous receptor signaling, with no discernible adaptation to more sustained proximal signaling. We were able to model the experimental kinetics as a simple linear chain of reversible reaction, with the most accurate fit as a sequence of five reactions (Figures 2L and S1I). From this we could estimate that it takes ~8 seconds for the information of signal cessation to propagate to the Ca^2+^ channels (1% decrease in output) and the effective half-life of proximal signal within this part of the network to be 29 seconds (95% CI: 27.4 – 30.5 s).

Overall, the implication of this dataset is that the opposing reactions along the signaling network from the receptor activation to the Ca^2+^ stores in T cells are efficient at dissipating proximal signaling once the stimulus has been removed, presenting very limited signal persistence at this level.

### ERK activation remains sensitive to the state of proximal signaling

The previous results suggested that continuous signaling from the receptor is required to maintain the intracellular fluxing of Ca^2+^ ions. We were therefore keen to understand whether this result held true for steps more distal from receptor activation in the T cell signaling network. A well-studied part of this pathway is the activation of extracellular signal-regulated kinase (ERK) by dual phosphorylation at Thr^102^ and Tyr^104^ (Roskoski, 2012). ERK is a classical member of the mitogen-activated protein kinase (MAPK) family and previous studies have suggested that ERK activation in T cells is digital (Altan-Bonnet and Germain, 2005; Das et al., 2009). We therefore wanted to use our OptoCAR system to investigate the dynamics of how the activating phosphorylation of ERK is modulated by the disruption of upstream signaling. Conjugates between OptoCAR-T cells and Raji-FRB^Ex^ cells were prepared in the absence of the dimerizer and placed in the blue light illuminator as used for the Ca^2+^ flux experiments. The dimerizer was then introduced and sample aliquots taken at defined timepoints by rapid fixation over a period of 30 minutes, with or without illumination. Cells were then stained for doubly phosphorylated ERK (ppERK) and the signal from conjugated cells was measured by flow cytometry.

Carrying out the experiment with the OptoCAR in the dark (signal-competent) state led to a rapid initial phosphorylation of ERK, with ppERK detectable within one minute before plateauing around 10 minutes (Figures 3A and 3D). The distributions of ppERK intensities over time show a clear all-or-nothing response to stimulation (Figure 3A), as previously reported (Altan-Bonnet and Germain, 2005). We were confident that this was the physiological response to receptor signaling because activation was synchronized by dimerizer addition and we specifically gated on OptoCAR-T cells conjugated to ligand-presenting cells. We then repeated the experiment but now illuminated the conjugates to disrupt receptor signaling after 5 minutes and continued to measure the distribution of ppERK staining. Illumination caused a rapid decrease in ppERK staining, which was detectable within 30 seconds and continued to decrease to baseline levels (Figures 3B and 3E). We were able to extract the half-life of the ppERK modification on cessation of proximal signaling, which we measured to be 3.0 minutes (95% CI: 2.4 – 3.6 min).

**Figure 3.**
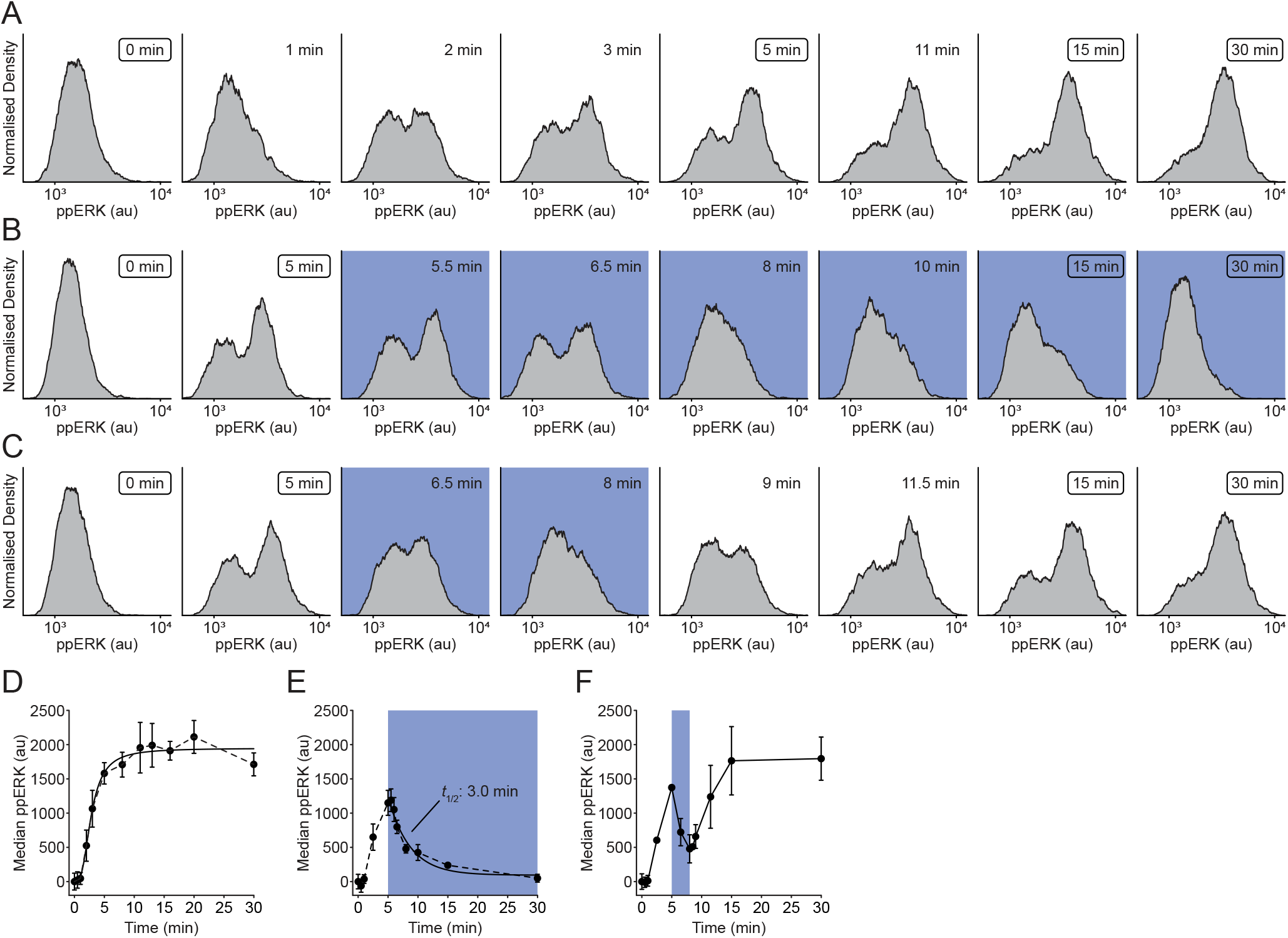
ERK activation remains sensitive to the state of proximal signaling. (A) OptoCAR-T cells were conjugated with ligand-presenting cells and receptor signaling was initiated with dimerizer addition at 0 minutes. Activated cells were then rapidly fixed at the timepoints specified in each panel and the distribution of doubly phosphorylated ERK (ppERK) was measured by flow cytometry, gated solely on cell conjugates. Boxed times indicate experimental points shared between all datasets. (B) An equivalent experiment was performed as in (A) but now illumination of the OptoCAR-T cell conjugates was initiated after 5 minutes for the remainder of the dataset. (C) An equivalent experiment was performed as in (A) but now illumination of the OptoCAR-T cell conjugates was initiated after 5 minutes for 3 minutes, before reverting to the dark (signaling competent) state of the OptoCAR for the remainder of the dataset. (D) Median ppERK levels were quantified from the datasets represented in (A) and plotted over time. Error bars show mean ± SEM (n=3). (E) Median ppERK levels were quantified from the datasets represented in (B) and plotted over time. Indicated half-life was calculated from the time taken for the median ppERK to decrease to 50% of maximum. Error bars show mean ± SEM (n=3). (F) Median ppERK levels were quantified from the datasets represented in (C) and plotted over time. Error bars show mean ± SEM (n=3).

Having shown that OptoCAR-mediated signaling was capable of activating ERK in a light-dependent manner, we wanted to investigate whether the observed inhibition of proximal signaling was reversible as we had found for Ca^2+^ fluxing (Figure 2E). After initiating T cell signaling with the dimerizer, we pulsed the cell conjugates with light after 5 minutes for 3 minutes before returning them to the signaling-competent state. As anticipated, ERK phosphorylation resumed quickly after resumption of receptor signaling at a rate that was equivalent to that observed prior to illumination (Figures 3C and 3F). Importantly, ppERK levels increased from the previous point immediately after reverting to the dark state, demonstrating that signal persistence within the network can lead to a more rapid increase in ppERK. However, the measured half-life of 3 minutes for ppERK decay after cessation of proximal signaling implies that any biochemical ‘memory’ at this part of the signaling network would completely dissipate by ~10 minutes.

### Activation of FOS transcription factor remains dependent on proximal signaling

The preceding dataset showed that the MAPK pathway remained rapidly reversible on termination of receptor signaling albeit with more persistence compared to the Ca^2+^ flux dynamics. We reasoned that this reversibility may extend to transcriptional activation within the nucleus. Many transcription factors (TFs) require phosphorylation to become active and enhance gene expression. This is true for the AP-1 family of leucine zipper TFs, typified by the FOS/JUN heterodimer (Karin et al., 1997). In T cells, FOS protein is rapidly expressed on activation as an immediate early gene (Clark et al., 2011; Ullman et al., 1990) and phosphorylation increases its TF activity to function, in conjunction with JUN, to upregulate expression of many genes (Karin et al., 1997).

We investigated the dependence of FOS phosphorylation on upstream signaling by conjugating the OptoCAR-T cells with ligand-presenting cells as normal in the presence of the dimerizer. We first measured the kinetics of *FOS* expression and its phosphorylation in the dark state to be sure that OptoCAR signaling could drive the activation of nuclear TFs. OptoCAR-T cells were activated for a given period before being rapidly lysed and samples subjected to fluorescent Western blot analysis. Both the phosphorylation of FOS at Ser^32^ (Figures 4A and S2A) and the total levels of FOS protein (Figures 4B and S2B) were quantified, with the former modification known to correlate with increased FOS TF nuclear localization and stability (Sasaki et al., 2006). We found that FOS protein was undetectable prior to activation in T cells but readily induced within 30 minutes (Figures 4B and 4C), and phosphorylation at Ser^32^ was detectable within 60 minutes (Figures 4A and 4D). Because neither the OptoCAR-T cells nor the Raji-FRB^Ex^ cells expressed FOS prior to activation, we could be confident that the detected protein arose solely from T cell activation. Total FOS protein peaked 2 hours after activation before decreasing (Figure 4C). However, quantifying the abundance of the Ser^32^ phosphorylated state as a fraction of total FOS showed that the nuclear-localized version continued to accumulate with increased signaling duration (Figure 4D). This effect was not due to the fraction of cells that were conjugated, which remained stable 60 minutes after dimerizer addition (Figure S2C).

**Figure 4.**
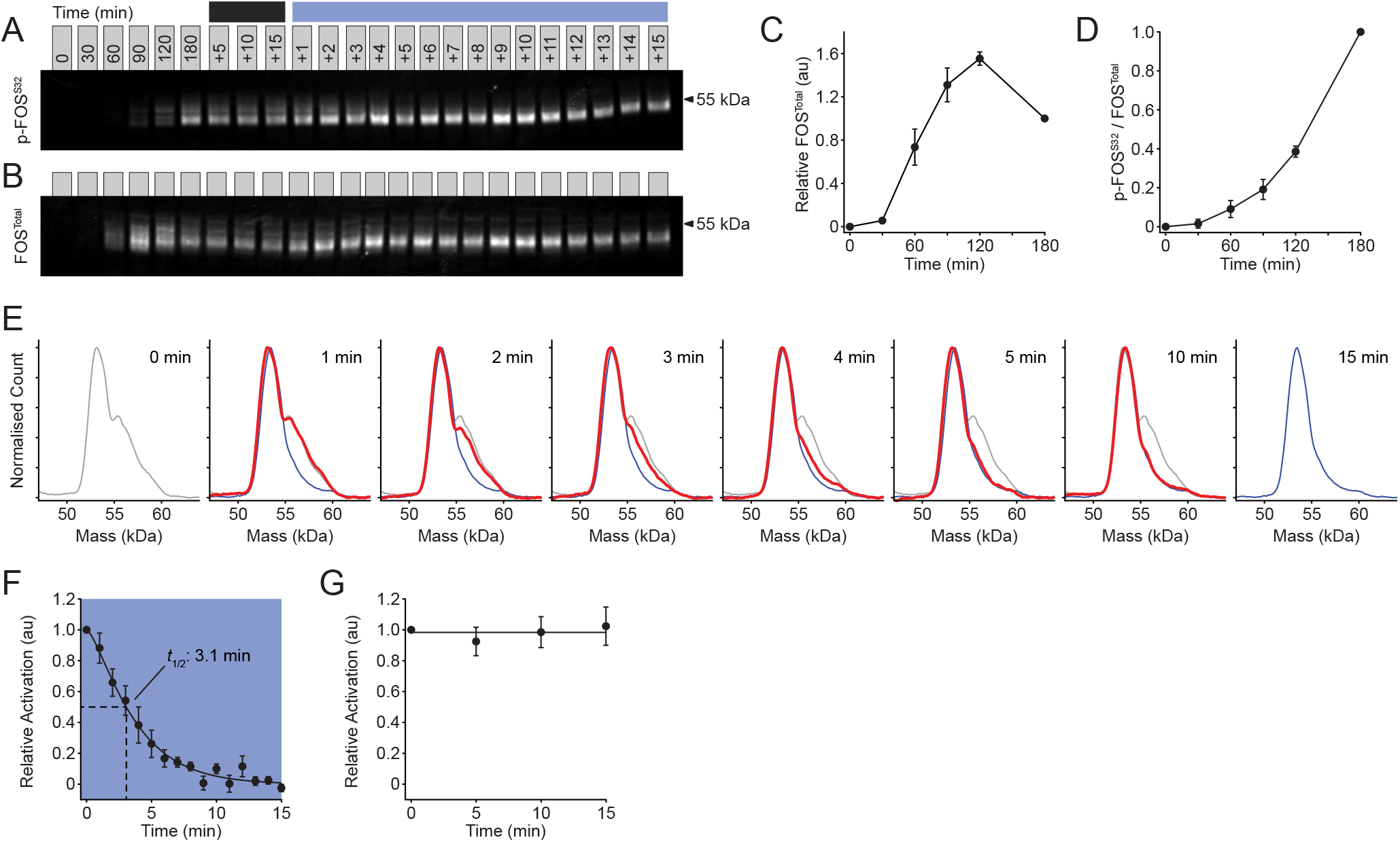
Activation of FOS transcription factor remains dependent on proximal signaling. (A) Representative fluorescent Western blot showing the dynamics of FOS phosphorylation (p-FOS^S32^) on light-modulated control. Conjugated OptoCAR-T cells were stimulated for a defined period (denoted above blots) before either being left in the dark (dark line) or illuminated (blue line). Full blot image with total protein normalization control shown in Figure S2A. (B) Representative fluorescent Western blot showing the dynamics of FOS expression (FOS^Total^) on light-modulated control, from the same dataset as in (B). Full blot image with total protein normalization control shown in Figure S2B. (C) Quantification of FOS expression detected by fluorescent Western blots at different timepoints prior to illumination. Datasets were normalized to FOS level at 3 hours. Error bars show mean ± SEM (n=3). (D) Quantification of the magnitude of phosphorylated FOS (p-FOS^S32^) as a fraction of total FOS, detected by fluorescent Western blots at different timepoints prior to illumination. Datasets were normalized to relative phosphorylated FOS level at 3 hours. Error bars show mean ± SEM (n=3). (E) Distributions of phosphorylated FOS molecular mass at different timepoints after illuminating OptoCAR-T cell conjugates to disrupt signaling, quantified from blot in (A). The distributions on initial illumination (0 min) and end of measurement (15 min) are shown as grey and blue lines, respectively. Distributions in other plots are shown as red lines that vary between these extremes. (F) The fraction of higher molecular weight phosphorylated FOS was quantified from the fluorescent Western blots and plotted over the time after illuminating the OptoCAR-T cell conjugates. Datasets are normalized between the values at 0 and 15 minutes. Indicated half-life was calculated from the time taken for the activated fraction to decrease to 50% of maximum. Error bars show mean ± SEM (n=3). (G) The fraction of higher molecular weight phosphorylated FOS was quantified from the fluorescent Western blots and plotted over the time over equivalent period to (F) but remaining the dark. Error bars show mean ± SEM (n=3). See also Figure S2.

Next, we investigated how light-mediated disruption of the OptoCAR in activated T cells controlled the fraction of active c-FOS TF. The primed active form of FOS runs as a higher molecular weight band by electrophoresis, compared to the unphosphorylated form (Murphy et al., 2002) and we used this feature to measure the fraction of functional FOS in our Western datasets (Figure 4A). After 3 hours of continuous stimulation in the dark, OptoCAR-T cell conjugates were illuminated using the OptoPlate described in the next section and samples taken at defined timepoints over 15 minutes. The disruption of OptoCAR-mediated signaling caused a readily detectable loss of the higher molecular weight phospho-FOS band (Figures 4A and 4E). Plotting the relative abundance of the activated form with time showed that essentially no higher molecular weight FOS TF could be detected after 10 minutes (Figure 4F). Fitting the individual datasets allowed us to quantify the half-life decay of FOS activity on cessation of proximal signaling to be 3.1 minutes (95% CI: 0.7 – 5.5 min). To ensure that this rapid decrease in TF activity was directly due to illuminating the cells, we measured the active FOS fraction over the same 15-minute period with the conjugated cells remaining in the dark state. As expected, this showed no significant decrease in the activation state of FOS (Figure 4G).

Overall, we have shown that the activation state of an important TF required for T cell activation remained intimately constrained by the signaling potential of the cell surface receptor, and confirmed the results above that the reverse steps in the signaling pathway are very efficient.

### Short-term signal persistence can be directly observed in gene expression output

The weak persistence of active FOS on disruption of receptor signaling agreed well with the equivalent dynamics of ppERK dephosphorylation, suggesting that much of the intracellular signaling network remains intimately connected to the functional state of the upstream receptor. This conclusion would suggest that any persistent state within the network could only have influence on the minutes timescale. However, it is possible that this short-term biochemical memory could nonetheless manifest in a more potent output response by integrating temporally discrete signaling pulses into more efficient gene expression. We therefore wanted to leverage the power of our optogenetic approach to directly test whether this was the case.

Receptor-mediated transcription factor activation should lead to *de novo* gene expression. To confirm that this was also true for OptoCAR-mediated signaling, we measured two functional outputs of cell activation (Figure 1A). The first output was from the functional activity of NFATc1-mediated GFP expression. NFATc1 (NFAT) is a TF that normally combines with AP-1 TFs to drive expression of key proteins, including the cytokine interleukin-2 that is critical for T cell activation. The second output is the increased expression of CD69 at the cell surface. We conjugated the OptoCAR-T cells as for the previous experiments whilst either providing continuous illumination or keeping them in the dark (signal-competent) state, and measured NFAT-mediated GFP expression and CD69 upregulation. As expected, OptoCAR-T cells activated in the dark for 24 hours showed potent upregulation of both functional outputs, but both responses were essentially undetectable under constant illumination during activation (Figures S3A and S3B).

Having shown that the OptoCAR could drive efficient gene expression, we next investigated how the duration of receptor signaling influenced the magnitude of the downstream outputs. This was accomplished by maintaining the conjugated cells in the dark to drive signaling for a defined period before disrupting OptoCAR signaling by illumination for the remaining time of a 24-hour experiment, before the downstream responses were measured. This experiment is therefore not a time course but a measure of the impact of signal duration on overall output. To perform these experiments, we employed a modified version of an ‘OptoPlate’ that is capable of independently illuminating the wells of a 96-well plate with control over both the duration and intensity of light pulses (Bugaj and Lim, 2019). We found that increased duration of signaling always resulted in greater output for both NFAT-mediated GFP expression (Figures 5A, 5B and S3C) and CD69 upregulation (Figures 5C, 5D and S3D), demonstrating that our system was not saturating at any point during the 24 hours of activation.

**Figure 5.**
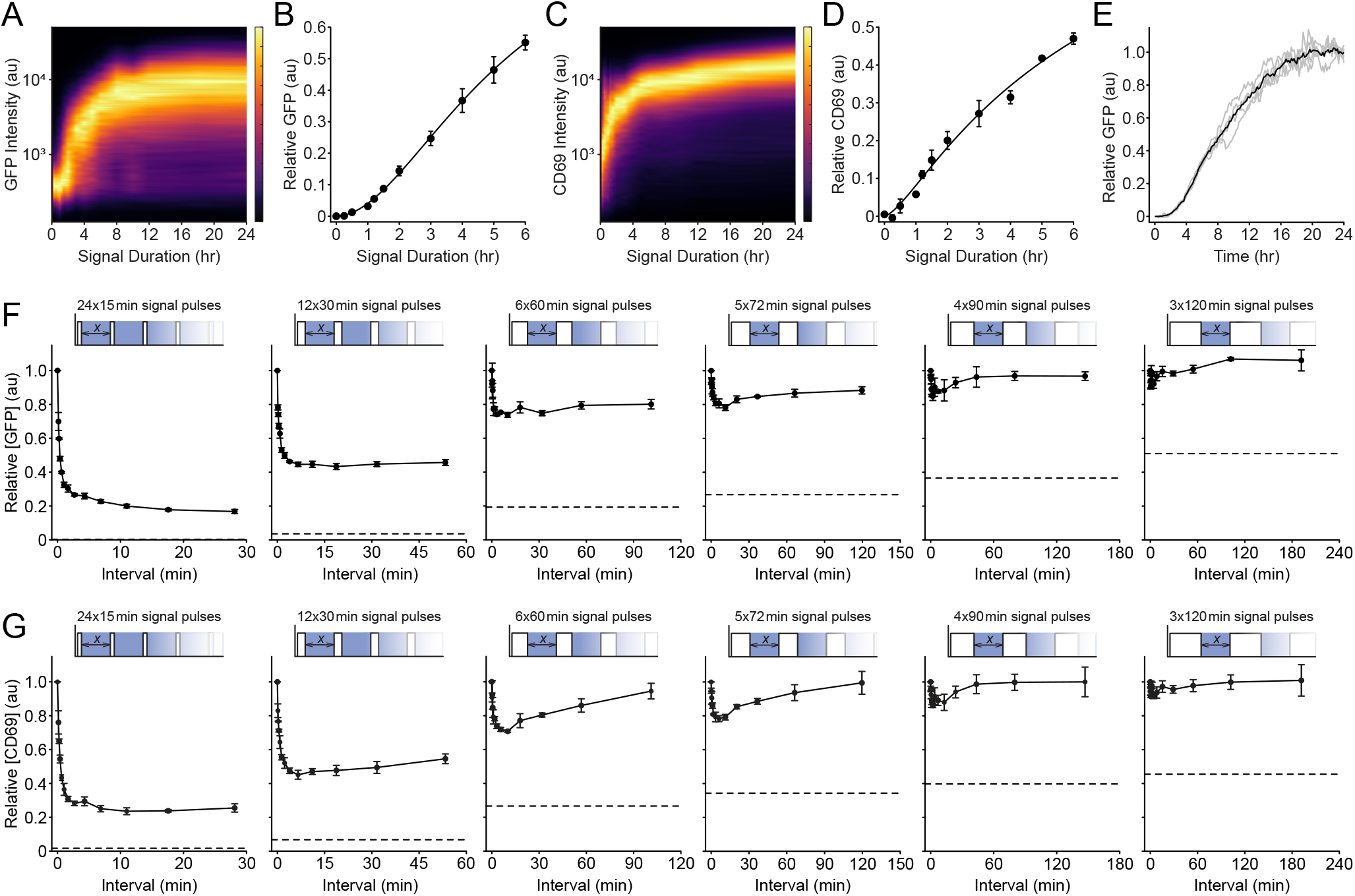
Short-term signal persistence can be directly observed in gene expression output. (A) Density plot of NFAT-mediated GFP expression as a function of varying the length of a single stimulus pulse. OptoCAR-T cell conjugates were activated in the dark (signaling competent) state for a defined period before light-mediated disruption of signaling for the remainder of a 24-hour period; cells were all assayed after this time. Plot is composed from 24 individual histograms, with a subset shown in Figure S3E. (B)Plot of geometric mean of GFP expression in (A) over a 6-hour range of signal pulses. The complete dataset is normalized to output at 24 hours, which is shown in Figure S3C. Error bars show mean ± SEM (n=3). (C) Density plot of CD69 upregulation as a function of varying the length of a single stimulus pulse, from same dataset as in (A). Subset of histograms are shown in Figure S3F. (D) Plot of geometric mean of CD69 upregulation in (C) over a 6-hour range of signal pulses. The complete dataset is normalized to output at 24 hours, which is shown in Figure S3D. Error bars show mean ± SEM (n=3). (E) The mean of NFAT-mediated GFP expression from OptoCAR-T cell conjugates over time, acquired by confocal microscopy. Individual traces from 5 stitched regions of experimental images are shown in grey lines, with mean time course in black. (F) Series of experiments showing how NFAT-mediated GFP expression is modulated by pulsatile trains of signaling. A combined signaling period of 6 hours was broken into pulses ranging from 15 minutes to 2 hours and the refractory period between these pulses varied (x-axis), shown in the schematic above each plot. Geometric mean of GFP intensity is shown plotted as a function of inter-pulse interval, normalized to continuous 6-hour output. Dotted line indicates output from an equivalent single pulse of activation. Error bars show mean ± SEM (n=3). (G) Series of experiments showing how CD69 upregulation is modulated by pulsatile trains of signaling, from same dataset as in (F). Error bars show mean ± SEM (n=3). See also Figure S3.

We also used live-cell confocal microscopy to measure the kinetics of NFAT-mediated GFP expression within cell conjugates. It took approximately two hours for GFP fluorescence to become detectable (Figure 5E), which would be expected from the combined requirements for maturation of the fluorophore as well as *de novo* gene expression. Almost all cells remained engaged with a ligand-presenting cell throughout the experiment, with dissociation invariably linked to the recent engagement of an alternative presenting cell (Movie S2).

The single-cell distributions of the outputs from flow cytometry suggested that the longer signaling duration increased the fraction of cells that responded rather than a uniform increase in the outputs (Figures S3E and S3F). We explored this response in more detail by continuously illuminating cells in each well of the 96 well plate over 24 hours but modulated the light intensity to directly titrate the input ‘strength’ emanating from the OptoCAR and quantified the cell’s output response to this linear input (Figure S4A). For both NFAT-mediated GFP expression (Figure S4B and Movie S3) and CD69 upregulation (Figure S4C and Movie S3) there was an initial regime under low illumination where the maximal output decreased in a graded response to decreasing signal intensity. However, at a certain threshold of decreased signaling driven by increased illumination, the output became digital in nature, with the activated fraction of cells being dependent on signaling intensity. A higher resolution dataset of this bistability in NFAT-mediated GFP expression with graded input provided a much clearer representation of the digital nature of this transition (Figure S4D and Movie S4) and suggested that some minimal level of signaling output must be reached before a cell becomes activated.

The duration of the minimal pulse that caused a detectable response was at least 30 minutes for both outputs, with CD69 upregulation showing the clearest response under brief stimulation (Figures 5B and 5D). As explained above, if any persistent signal remained within the signaling network then multiple pulses of these short periods of stimulation should drive a more robust downstream cellular output. Thus, we separated a 6-hour total stimulus into a sequence of multiple pulses and varied the interval between them. We used signal pulses ranging from 15 to 120 minutes in duration and measured the downstream outputs of T cell activation after 24 hours. For example, 5 ‘dark’ signal pulses of 72 minutes were applied to conjugated OptoCAR-T cells with the interval between the pulses (i.e., illuminating the cells) ranging from 0 to 120 minutes. If there was no signal persistence within the network at the end of each pulse, then the cell’s output should be independent of the interval length; conversely, any integration with previous pulses would show a dependence on interval duration.

By encoding these pulsatile stimulation profiles onto conjugated T cells, we could readily observe an interval-dependent decrease in both NFAT-mediated GFP expression (Figure 5F) and CD69 upregulation (Figure 5G). Pertinently, this dependence decayed rapidly within 10 minutes for all stimulus periods (Figures S3G and S3H), in excellent agreement with the kinetics of loss of active ERK (Figure 3E) and FOS (Figure 4F) TFs on signal cessation. This dataset is strong evidence that persistence of residual signaling within the intracellular signaling network can be directly observed at the level of transcriptional output, but that it rapidly dissipates.

### Gene transcription is rapidly abolished after disruption of receptor signaling

While the overall output measured with pulsatile stimulation decreased rapidly when the inter-pulse interval increased over 10 minutes, it plateaued to a level significantly higher than that measured for an equivalent length single pulse. This was especially evident for the 15-minute pulse datasets, where a single pulse gave an undetectable response, but repeated pulses became detectable. This implied that an additional intermediate within the signaling network could persist for longer between pulses. To explore whether sustained promoter activity was responsible for this, we used reverse transcription quantitative PCR (RT-qPCR) to measure the relative mRNA concentration of several upregulated genes during OptoCAR-T cell activation over 3 hours, before illumination and collection of samples over a further 60 minutes. Samples at equivalent time points were also prepared under continued dark conditions.

The mRNA from genes with distinct TF requirements in Jurkat T cells were quantified: CD69 is predominantly activated by AP-1 (Castellanos et al., 1997), efficient IL2 expression is dependent on CD28-mediated NFAT and AP-1 TFs (Shapiro et al., 1997; Spolski et al., 2018), CXCL8/IL8 transcription requires NF-ĸB activity (Hoffmann et al., 2002; Kunsch and Rosen, 1993) and FOS transcription is dependent on Elk-1 (Cavigelli et al., 1995). The genes investigated presented a range of expression profiles, with *FOS* showing the hallmarks of an immediate-early gene as expected, and *IL2* a delayed response likely due to combining signals from the OptoCAR and CD28 costimulatory receptor (Figure 6A). However, all mRNA transcripts showed a pronounced decrease on illumination that approached unstimulated levels over 60 minutes (Figures 6A and 6B). While there appeared to be continued gene transcription for CD69 and IL2 after receptor input termination (Figure 6B), correcting for the underlying trajectory of mRNA expression by taking the ratio between the paired illuminated and dark samples, we found that mRNA levels started to decrease for all genes within 5 minutes of illumination (Figure 6C). This result implied that nascent mRNA production is halted at an equivalent timescale to inhibition of TF activity, with the observed decay due to the mRNA degradation pathways, assumed to be independent of the active state of the promoter. Thus, these decay rates with an approximate half-lives of 15-30 minutes are the limiting bound on how long signals are sustained within the network and would explain how single undetectable pulses can be combined into a detectable output.

**Figure 6.**
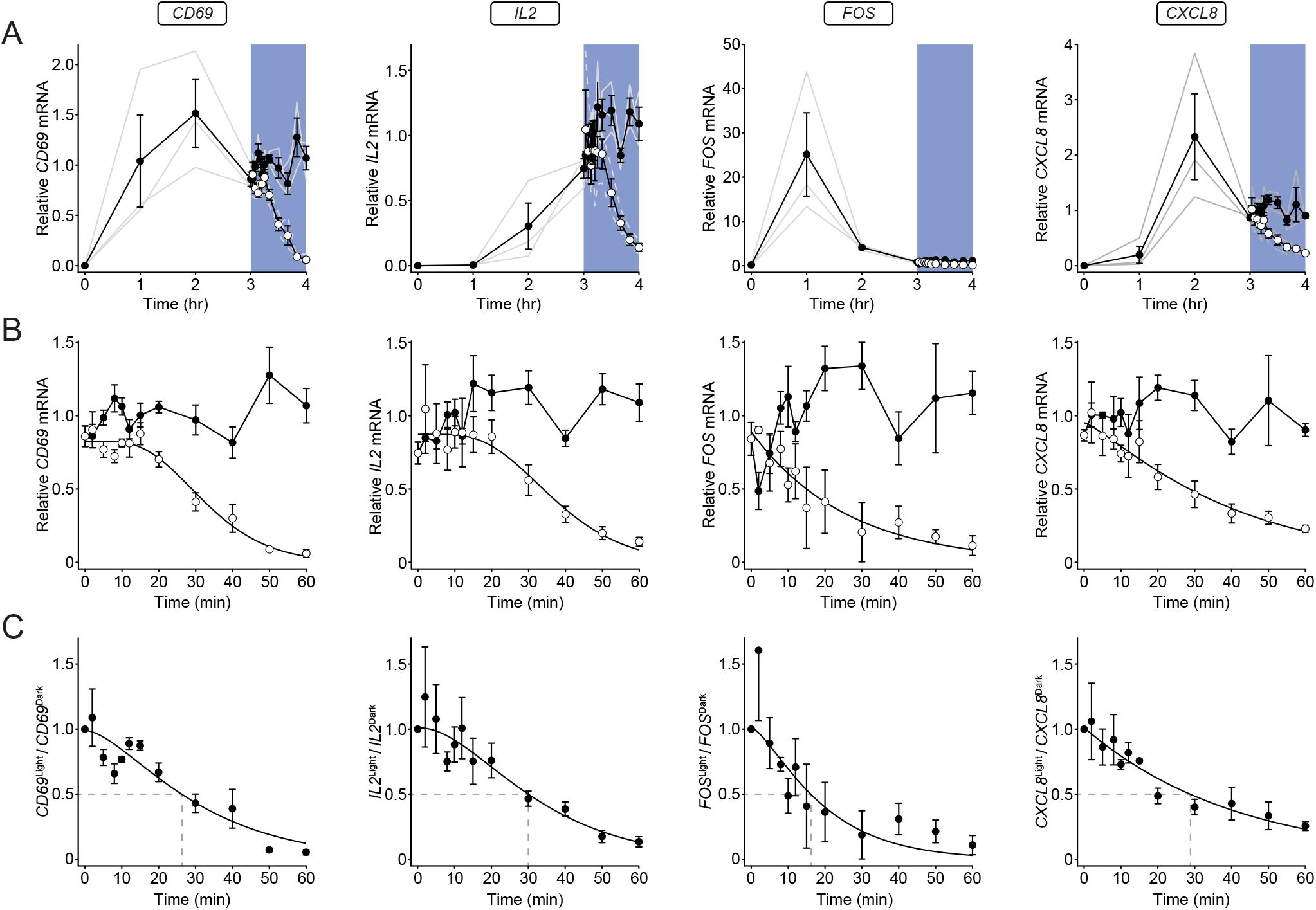
Gene transcription is rapidly disrupted on cessation of OptoCAR signaling. (A) RT-qPCR analysis of relative mRNA levels for four representative genes following OptoCAR-mediated T cell activation. OptoCAR-T cell conjugates were activated in the dark (signaling competent) state for a defined period before light-mediated disruption of signaling after 3 hours (open circles) or maintained for a further 60 minutes in the dark (filled circles). Individual datasets are presented as gray lines, with error bars show mean ± SEM of 3 biological replicates. (B) Equivalent datasets expanded to show the effect of illumination on the relative mRNA level of designated genes. As above, datasets are normalized to the mean mRNA level for the dark state values over the 60 minute period. Error bars show mean ± SEM (n=3). (C) The ratio of relative mRNA in the light compared to dark state were calculated for each timepoint. An approximate half-life for mRNA decay is shown with dotted lines. Error bars show mean ± SEM (n=3).

### Pulsatile signaling enhances the output of a clinically relevant anti-CD19 CAR

A sustained and potent stimulus can drive negative feedback loops within signaling networks (Amit et al., 2007) and there is evidence that a pulsed signaling input might alleviate this signaling-induced negative functional state (Wilson et al., 2017). Excessive or prolonged receptor signaling can drive primary T cells into a dysfunctional or ‘exhausted’ state (Blank et al., 2019) and is commonly found with chimeric antigen receptors (CARs) currently used clinically (CAR-T therapy) to treat blood cancer patients (Wu et al., 2020). We therefore wanted to use our optically controlled receptor strategy to test whether pulsed CAR input might drive more effective T cell activation. To this end, we engineered optical control into a clinically relevant anti-CD19 CAR (Porter et al., 2015), forming OptoCAR^CD19^ (Figure 7A) and transduced primary CD4^+^ T cells with this construct. We first confirmed that this modification to the CAR maintained its function by conjugating these OptoCAR^CD19^ expressing T cells with a CD19^+^ B cell line and measuring expression of CD137 (4-1BB), which is a robust marker of T cell activation (Figure 7B). As expected, we found that continuous blue light illumination completely abrogated CD137 expression (Figure 7B).

**Figure 7.**
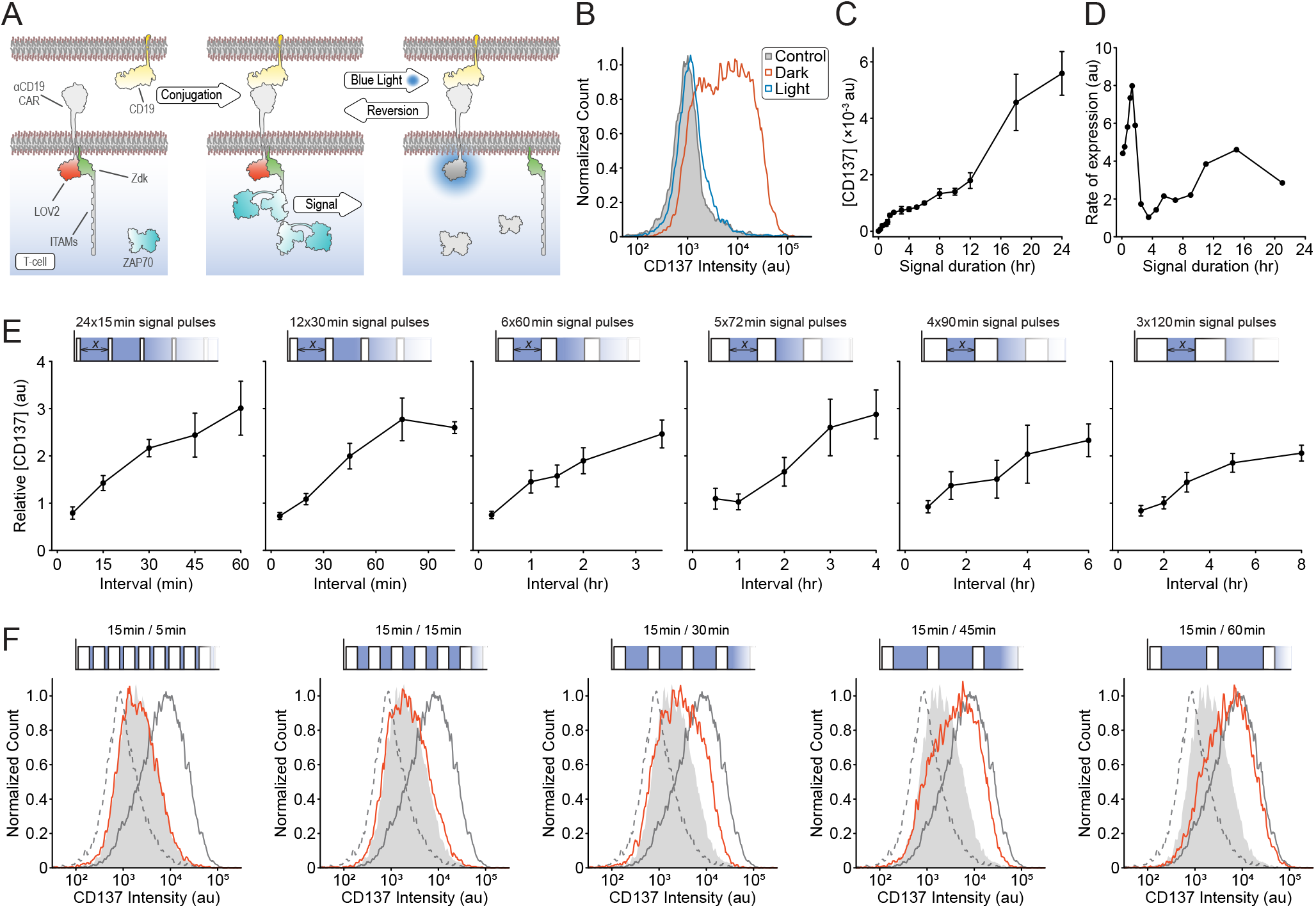
anti-CD19 CAR-T cell activation can be enhanced by pulsatile signaling. (A) Schematic showing the anti-CD19 OptoCAR engineered to be light-responsive. Binding of the OptoCAR^CD19^ expressed in T cells to CD19-expressing B cells drives signaling through an equivalent intracellular sequence used in the OptoCAR, where illumination with blue light causes the reversible disruption of receptor signaling. (B) Stimulating primary CD4^+^ T cells expressing OptoCAR^CD19^ over 24 hours in the dark drives expression of CD137 (4-1BB), a robust activation marker. Continuous illumination completely abolishes this to CD137 levels equivalent to unconjugated T cells. (C) Plot of geometric mean of CD137 expression over a 24-hour range of signal pulses. The complete dataset is normalized to output at 6 hours. Sample illumination was initiated at the end of the pulse and CD137 expression measured for all samples after 24 hours. Error bars show mean ± SEM (n=3). (D) The approximate expression rate of CD137 was determined by taking the differential of the data in (C), with a clearly observed maximum around 60 minutes. (E) Series of experiments showing how CD137 expression is modulated by pulsatile trains of signaling in primary CD4^+^ T cells activated through OptoCAR^CD19^. A combined signaling period of 6 hours was broken into pulses ranging from 15 minutes to 2 hours and the refractory period between these pulses varied (x-axis), shown in the schematic above each plot. Geometric mean of CD137 intensity is shown plotted as a function of inter-pulse interval, normalized to continuous 6-hour output. Error bars show mean ± SEM (n=3). (F) Representative dataset from (E), demonstrating how pulsatile stimulation of OptoCARCD19-T cells drives more efficient activation. Dotted and gray histograms show CD137 expression in resting cells and 24-hour activation, respectively. The filled histogram shows continuous 6-hour stimulation, whereas the red histogram shows equivalent pulsed activation over 24 hours driving substantially more potent activation.

We next varied the duration of a single pulse through the OptoCAR^CD19^ and measured CD137 expression after 24 hours. In contrast to the equivalent experiment with the chemically-inducible OptoCAR (Figures S3C and S3D), while there was an initial correlation between signal duration and cellular output, this plateaued after ~2 hours with only a slow increase in output for pulses up to ~12 hours (Figure 7C). Estimating the rate of activation from this dataset showed a maximum around a 60-minute pulse (Figure 7D). We hypothesized that this outcome was due to signaling-induced feedback inhibition of CD137 expression and that by using short pulsatile bursts of signaling we might be able to alleviate this inhibition. As before, we split a single 6-hour stimulus into pulses ranging from 15 minutes to 2 hours and measured CD137 expression after 24 hours (Figure 7E). The response to pulsed signaling was very pronounced, with 15-minute pulses leading to an approximately three-fold increase in output when separated by up to 60 minutes that approached CD137 expression found after a 24-hour period (Figure 7F).

## DISCUSSION

In this study, we have described an engineered antigen receptor that provides optical control over intracellular signaling in T cells whilst conjugated with antigen presenting cells. We have used this new tool to directly interrogate the intracellular network’s capacity for antigen receptor signal integration by quantifying the persistence of stimuli at various points within this network. The main finding from this work is that at all parts of the intracellular signaling network tested, disruption of antigen receptor signaling caused essentially all residual information within the network to dissipate within 10-15 minutes. An important corollary of these conclusions is that the opposing reactions within the signaling network that seek to re-establish the basal level of signaling must be very efficient and potent, with continuous signaling flux required to overcome them. Nevertheless, we could directly observe the persistence of this biochemical signal within the network at the gene expression level using pulsatile trains of signaling. We also demonstrated that sustained proximal signaling is required to maintain transcription factors in an active state, and hence continued gene expression, with the decay of mRNA being the longest-lived signaling intermediate having approximate half-lives between 15-30 minutes for the transcripts measured here. We then demonstrated that this limited signal persistence could be exploited to minimize the induction of inhibitory feedback when T cells are over-stimulated by a clinically relevant anti-CD19 CAR through pulsatile signaling.

Our results imply that T cells are not capable of directly integrating TCR signals over multiple APC interactions beyond a short temporal window. However, there is good evidence that T cells do accumulate the output from gene expression over multiple interactions (Clark et al., 2011; Faroudi et al., 2003; Munitic et al., 2005), suggesting that a threshold of protein expression must exist beyond which the specification of T cell function occurs. By titrating the magnitude of the signal intensity emanating from the OptoCAR using graded illumination, we were able to demonstrate that a minimal level of signaling is required to drive downstream output in a digital manner, in agreement with this. The study from Bousso and colleagues (Clark et al., 2011) suggested that the accumulation of activated FOS was a potential mechanism for integrating multiple T cell signaling events, with phospho-FOS^T325^ increasing even when receptor signaling is disrupted. However, this result does not fit with the rapid loss of this modification when ERK activity is inhibited (Murphy et al., 2002), in agreement with our finding that FOS activity is rapidly lost on cessation of signaling (Figure 4F). This discrepancy may lie in how activated FOS was detected or the use of antibody-coated beads to activate the T cells.

Previous studies have investigated how transient inhibition of TCR signaling affects the level of Ca^2+^ fluxing and the rate of its decline on signal disruption (Valitutti et al., 1995; Varma et al., 2006; Yousefi et al., 2019). These reports found a strong dependence on proximal TCR signaling to maintain the increased level of Ca^2+^ ions in agreement with our findings. However, we measured a far more rapid decrease in this readout (Figure 2L), which we attribute to the way we disrupt receptor signaling being more direct and efficient. The rapid decrease in phosphorylated ERK on disruption of signaling that we observed (Figure 3E) has also been found in another recent optogenetic study in NIH3T3 cells (Bugaj et al., 2018), suggesting that the results we have found are likely to be more generally applicable beyond T cell signaling. Our finding that the active state of the transcription factor FOS, like ERK, required continuous proximal signaling implies that only their downstream output of increased gene expression can constitute a significant persistent state of previous signaling. This means that the encoding of signaling dynamics must be at this level too and previous studies have provided evidence for this result (Clark et al., 2011; Locasale, 2007; Marangoni et al., 2013; Murphy et al., 2002).

There have been several other studies employing optogenetics to investigate how signaling dynamics influence downstream cellular activation, which have led to some exciting new results (Bugaj et al., 2018; Graziano et al., 2017; Toettcher et al., 2013; Wilson et al., 2017). These studies invariably used the light-mediated translocation of a constitutively active enzyme to the plasma membrane as the means to control downstream signaling. While very effective, this approach ‘short-circuits’ signaling from the upstream receptor, potentially bypassing key parts of the network and removing any spatio-temporal information that might be encoded in physiological receptor activation within cell conjugates. In our approach, light controls the very initiation of receptor signal transduction without altering the network architecture, with receptor activation occurring within the physiological context of a T cell/presenting cell conjugate interface. We believe this approach provides the most realistic control over receptor activation, without disrupting signaling from other cell-surface receptors that might modulate the TCR input, such as co-stimulation through CD28 engagement. We have also investigated the endogenous signaling components rather than over-expressing protein sensors, which should most faithfully report the dynamics of the network.

The OptoCAR we have developed in this work contains three ITAM signaling motifs. While capable of initiating very efficient downstream signaling, it cannot be expected to fully replicate all the aspects of the complete TCR complex. We have previously shown that the orthogonal receptor/ligand pair used in the OptoCAR can drive equivalent segregation of the phosphatase CD45 that is thought to initiate receptor signaling as observed for the TCR (James and Vale, 2012) as well driving downstream signaling (James, 2018). We have also shown that the increased number of ITAMs present on the TCR allows it to be highly efficient at transducing ligand binding into a signal even at low receptor occupancy (James, 2018). However, for all the experiments performed in this work we have worked in a high occupancy regime by saturating with the dimerizer, so we expect that the OptoCAR is provided a broadly equivalent stimulus to that expected from the TCR itself.

We have used the Jurkat cell line for most of our assays, which is a well-used model for T cell studies (Abraham and Weiss, 2004) but cannot entirely replicate the functions of primary CD4^+^ T cells (Bartelt et al., 2009). However, the known mutation in Jurkat cells in PI3K signaling due to PTEN inactivation (Shan et al., 2000) would not be expected to have any direct effect on the dynamics of the signaling pathways investigated in this study. We are therefore confident that the conclusions we have drawn from our datasets are likely to generalize to other receptor signaling networks that depend on the same transduction pathways.

## Supporting information

Supplementary Figures (Combined)

## ACKNOWLEDGMENTS

We are very grateful to the mechanical and electronic workshops at the MRC-LMB for assistance with construction of the illumination devices used in this study, and to Klaus Hahn and Lukasz Bugaj for sharing reagents that were invaluable to this work. We thank members of the James lab for commenting on the manuscript. This work was supported by the Wellcome Trust (grants 099966/Z/12/Z to J.R.J. and 102195/Z/13/Z to M.J.H) and Warwick Medical School. M.F acknowledges funding from the University of Warwick, the EPSRC & BBSRC Centre for Doctoral Training in Synthetic Biology (grant EP/L016494/1). We acknowledge equipment access, training and support made available by the Research Technology Facility of the Warwick Integrative Synthetic Biology centre (WISB), which received funding from EPSRC and BBSRC (BB/M017982/1).

## AUTHOR CONTRIBUTIONS

Conceptualization, M.J.H. and J.R.J.; Methodology, M.J.H., M.F. and J.R.J.; Investigation, M.J.H., M.F. and J.R.J.; Writing – Original Draft, J.R.J.; Writing – Review & Editing, M.J.H., M.F. and J.R.J.; Visualization, J.R.J.; Supervision, J.R.J.; Funding Acquisition, J.R.J.

## STAR★METHODS

### KEY RESOURCES TABLE

| REAGENT or RESOURCE | SOURCE | IDENTIFIER |
| --- | --- | --- |
| **Antibodies** |  |  |
| Mouse anti-human CD86 (Alexa Fluor 647) | Biolegend | Cat#: 305416 |
| Human TruStain FcX (FcR blocking) | Biolegend | Cat#: 422302 |
| Phospho-Erk1/2 (Thr<sup>202</sup>/Tyr<sup>204</sup>) mAb (Alexa Fluor 488) | CST | Cat#: 4374S |
| Rabbit anti-Phospho-FOS (Ser32) mAb | CST | Cat#: 5348S |
| Rabbit anti-human FOS mAb | CST | Cat#: 2250S |
| Mouse anti-human CD69 (Alexa Fluor 647) | Biolegend | Cat#: 310918 |
| Mouse anti-human CD3 (Alexa Fluor 647) | Biolegend | Cat#: 300322 |
| Mouse anti-human CD20 (Brilliant Violet 605) | Biolegend | Cat#: 302334 |
| Mouse anti-human CD137 (Brilliant Violet 421) | Biolegend | Cat#: 309820 |
| **Bacterial and Virus Strains** |  |  |
| Escherichia coli DH5alpha | ThermoFisher | Cat#: 18265017 |
| **Chemicals, Peptides, and Recombinant Proteins** |  |  |
| A/C Dimerizer (AP21967) | TaKaRa | Cat#: 635057 |
| Indo-1 LR (AM) | TEFLab | Cat#: TEF0145 |
| Pluronic F-127 | Sigma-Aldrich | Cat#: P2443 |
| Lenti-X Concentrator | TaKaRa | Cat#: 631231 |
| Perm Buffer III | BD | Cat#: 558050 |
| **Critical Commercial Assays** |  |  |
| RNeasy Plus Mini Kit | Qiagen | Cat#: 74134 |
| QIAshredder | Qiagen | Cat#: 79654 |
| SuperScript IV VILO Master Mix with ezDNase | Thermo Fisher | Cat#: 11756050 |
| TaqMan Advanced Fast Master Mix | Thermo Fisher | Cat#: 13428456 |
| *GAPDH* TaqMan Assay | Thermo Fisher | Hs02786624_g1 |
| *PGK1* TaqMan Assay | Thermo Fisher | Hs00943178_g1 |
| *CD69* TaqMan Assay | Thermo Fisher | Hs01119267_g1 |
| *FOS* TaqMan Assay | Thermo Fisher | Hs00934033_m1 |
| *IL2* TaqMan Assay | Thermo Fisher | Hs00174114_m1 |
| *CXCL8 (IL8)* TaqMan Assay | Thermo Fisher | Hs00174103_m1 |
| **Experimental Models: Cell Lines** |  |  |
| HEK-293T: Human adherent cell-line | ATCC | Cat#: CRL-11268 |
| J.NFAT: Derived from human Jurkat T-cell leukemia | (James, 2018) | N/A |
| Raji-FRB<sup>Ex</sup>: Derived from human B-cell lymphoma line | (James and Vale, 2012) | N/A |
| **Oligonucleotides** |  |  |
| For primer sequences, see Figure S5 | This work | N/A |
| **Recombinant DNA** |  |  |
| pCMVΔR8.91 | (Zufferey et al., 1997) | N/A |
| pMD2.G | Gift from Didier Trono | Addgene#: 12259 |
| pHR-OptoCAR-mScarlet | This work | N/A |
| pHR-iRFP670-CaaX | This work | N/A |

| Software and Algorithms |  |  |
| --- | --- | --- |
| FIJI (ImageJ): Image analysis | (Schindelin et al., 2012) | https://fiji.sc/ |
| MATLAB R2019a: Data processing | Mathworks | N/A |
| FlowJo V10.2: Data processing | FlowJo LLC | N/A |
| Excel 365 | Microsoft | N/A |
| Illustrator 2020 | Adobe | N/A |

### RESOURCE AVAILABILITY

#### Lead Contact

Further information and requests for resources, reagents and materials should be directed to and will be fulfilled by the Lead Contact, John R. James.

#### Materials Availability

All unique/stable reagents generated in this study are available from the Lead Contact with a completed Materials Transfer Agreement. Plasmids generated in this study will be deposited in Addgene.

#### Data and Code Availability

The MATLAB scripts used to process data during this study are available from the lead contact on reasonable request.

### EXPERIMENTAL MODEL AND SUBJECT DETAILS

#### Cell culture

HEK-293T cells were grown in Dulbecco’s modified Eagle’s medium supplemented with 10% fetal bovine serum (FBS) and antibiotics. Jurkat (J.NFAT) cells (James, 2018) and Raji-FRB^Ex^ cells (James and Vale, 2012) were grown in RPMI 1640 medium, supplemented with 10% FBS, 10 mM HEPES, and antibiotics. All cell lines were maintained at 37°C with 5% CO_2_ in a fully humidified incubator.

### METHOD DETAILS

#### OptoCAR vector construction and lentiviral transduction

To create the optically controllable chimeric antigen receptor termed OptoCAR in the main text (Figures S5A and S5B), we first amplified the wildtype LOV2 light-sensing domain from *Avena sativa* phototropin 1 using primers pr1/pr2 (Figure S5C). We then fused this part to the extracellular and transmembrane regions of our previous chemically-inducible CAR, FKBP-CD86-CD3ζ (James, 2018) by inserting it as a *Spe*I/*Not*I fragment, replacing the ζ-chain sequence. This FKBP-CD86-LOV2 construct was then excised as a *Mlu*I/*Not*I fragment and inserted into a vector (pHR-1G4) that can accommodate two genes separated by the P2A ribosomal skip sequence (James, 2018). The intracellular sequence of the bipartite OptoCAR was gene synthesized (LCK myristoylation sequence, Zdk, ζ-chain and mScarlet fluorophore; see Figure S5A) and fused to P2A and FKBP-CD86-LOV2 by overlap extension PCR using primers pr3-pr5 and pr1 (Figure S5C). This process also removed a *Spe*I site at the 3’ end of mScarlet to make it unique at the 5’ end of LOV2. The single contiguous open reading frame inserted into pHR-1G4 as an *Asi*SI/*Not*I fragment (Figure S5A).

To create LOV2 mutants for some of the OptoCAR experiments (C450G [TGC->GGC], I539E [ATT->GAA], V416L [GTC->CTC]), we used site-directed mutagenesis and cloned them as *Spe*I/*Not*I fragments into the OptoCAR vector. All vector sequences were confirmed by Sanger sequencing. For the variant of the OptoCAR that used a cytoplasmic intracellular sequence, the complete section of Zdk-CD3ζ-mScarlet was simply amplified without the myristoylation region (but now including a Kozak sequence) using pr6/pr7 (Figure S5C) and replaced the equivalent region in the normal OptoCAR vector as a *Asi*SI/*Bam*HI fragment.

The OptoCAR sequence was introduced into J.NFAT cells by lentiviral transduction in an equivalent manner described previously (James, 2018). This invariably led to cells that were uniformly positive and had a tight distribution of expression such that there was no need to sort or clone OptoCAR-T cells.

For the OptoCAR^CD19^ construct (Figures S5D and S5E), the extracellular and transmembrane region of anti-CD19 CAR was amplified as a *Mlu*I/*Spe*I fragment and replace the equivalent section in the OptoCAR vector.

#### Flow Cytometry

Cell samples to be analyzed by flow cytometry in standard 12×75 mm tubes were centrifuged at 800×*g* for 3 minutes, with pellets flick resuspended. Where required, primary antibodies were used at a working concentration of ~10 μg/mL, diluted in Flow wash buffer [2.5% (v/v) FBS, 0.1% (w/v) NaN_3_ in PBS, pH 7.4] and incubated on ice for at least 30 min, with regular agitation. For Raji B cells, endogenous Fc receptors were pre-blocked with Human TruStain FcX for 10 minutes at room temperature. Samples were washed with 3 mL Flow wash buffer and centrifuged as before. When the primary antibody was fluorescently conjugated, the pellet was resuspended and fixed in Flow fix buffer [1.6% (v/v) formaldehyde, 2% (w/v) glucose, 0.1% (w/v) NaN_3_ in PBS, pH 7.4). When a secondary labelled antibody was required, an equivalent staining protocol was used, with the secondary antibody at 20 μg/mL, prior to washing and fixing.

For staining in 96-well round-bottomed plates, samples were transferred into the plate using a multichannel pipette, ensuring cells were properly mixed by gentle pipetting up and down. The plate was then centrifuged at 800×*g* for 3 minutes and the supernatant was discarded. Cell pellets were resuspended in the residual media by pulsed vortexing. When required, primary antibodies at ~10 μg/mL, diluted in Flow wash buffer (100 μL/well) were added to all wells and incubated on ice for at least 30 min. Wells were then washed by adding 150 μL FACS wash and the plate was centrifuged at 800×*g* for 3 minutes. The supernatant was discarded, and the plate vortexed to resuspend the cell pellet in the residual solution. A second wash with 200 μL was performed similarly. Finally, 200 μL FACS fix buffer was then pipetted into the wells and the plate stored at 4°C until analyzed.

The flow cytometer was set up to ensure the most appropriate dynamic range between the negative and positive controls for each fluorescence channel and at least 30,000 gated cells were collected for each analysis. All data were collected on either a BD LSRII or Fortessa flow cytometer, with the latter instrument having an HTS unit to collect data from the 96-well plates.

#### Measurement of OptoCAR dissociation kinetics by microscopy

To quantify the rates at which the bipartite OptoCAR construct dissociated on illumination and reformed on return to the dark condition, we transiently transfected HEK-293T cells with the OptoCAR variant that had a cytoplasmic intracellular part, in combination with prenylated iRFP670 fluorophore (iRFP-CaaX) to demark the plasma membrane. Transfected cells were then adhered to fibronectin-coated imaging dishes and placed on a spinning-disk confocal microscope that was maintained at 37°C and 5% CO_2_. A 100× oil-immersed objective was used to image the OptoCAR construct (mScarlet: 561nm Ex; 607±18nm Em) and iRFP-CaaX (640nm Ex; 708±37nm Em) sequentially at 1 Hz. These fluorophores were utilized as imaging them would not have any influence on the conformational state of LOV2. To simultaneously image and illuminate cells, we placed a blue (450±25nm) filter in the condenser of the transmitted light path using a custom-designed holder and closed down the field diaphragm so only the cells in the field-of-view were illuminated. After 25 seconds of imaging, the transmitted white light LED was switched on to illuminate the sample region in focus for a further 25 seconds before turning the LED off. Imaging continued for a further 100 seconds.

Image stacks were subsequently analyzed in MATLAB. The two channels were first registered to optimally align the stacks before the cell regions of interest (background, cytoplasm, whole cell) were user-defined and both channels were background subtracted and bleach corrected. The iRFP-CaaX channel was then thresholded and Gaussian bandpass filtered to define a binary representation of the plasma membrane for each frame. This mask was then used to calculate the mean intensity of the OptoCAR intracellular section compared to the mean intensity from the cytoplasm for each frame.

#### Formation of cell conjugates

Many of the assays required conjugating J.NFAT cells expressing the OptoCAR with Raji-FRB^Ex^ cells in the absence of the dimerizer, prior to starting the experiment itself. Generally, a specified number of OptoCAR-T cells and Raji-FRB^Ex^ cells were first centrifuged separately at 800×*g* for 3 minutes and the cell pellet flick resuspended in residual medium. A small volume of fresh medium was then added to the OptoCAR-T cells before combining with the Raji-FRB^Ex^ cells, such that the cells were at high density at a 1:1.5–2 ratio. Usually, this cell mixture was then aliquoted into separate 1.5 mL Eppendorf tubes and centrifuged at 800×*g* for 1 minute to drive the cells into close apposition and incubated at 37°C for between 10 and 30 minutes. After this incubation, the cell pellets in each tube were gently resuspended by pipetting up and down, fresh medium was added to increase the volume and then used directly in the required downstream application.

#### Measurement of Ca^2+^ flux by flow cytometry

OptoCAR-T cells were labelled with the fluorescent Ca^2+^ indicator, Indo-1 AM by resuspending 5×10^5^ cells in 500 μL serum-free medium containing Indo-1 at a final concentration of 5 μg/mL, supplemented with 0.1% Pluronic F-127. The cells were loaded with Indo-1 at 37°C for 30 minutes. Labelled cells were then washed using 3 mL PBS (with Ca^2+^ and Mg^2+^) to remove any excess Indo-1 and flick resuspended in residual volume. Conjugates of Indo-1 loaded cells and Raji-FRB^Ex^ cells were prepared as described above and added to standard flow cytometry tubes ready for sample acquisition.

To optically modulate the conjugated cells whilst measuring the intracellular [Ca^2+^] on the flow cytometer, we custom-built a device that could illuminate the sample tube with a 455nm wavelength LED (royal blue) whilst inserted in the cytometer. We also included a thermostatically controlled heating block around the tube so that the desired temperature could be maintained during data acquisition. Conjugated cells were run on the flow cytometer for 60 seconds to define a baseline ratiometric signal of Indo-1 fluorescence (405±15 nm / 485±13 nm) excited at 355 nm. The sample tube was then removed and 2 μM dimerizer added to initiate signaling within the conjugates, which were gated on as iRFP713^+^/mTagBFP^+^ doublet events. Events were collected for varying periods depending on the type of experiment and when required, the LED illuminator was switched on manually at the appropriate time.

For data analysis, cytometry files were first gated in FlowJo before exporting CSV data files that were further processed in MATLAB. The Indo-1 ratio was calculated after background subtraction from the individual channels and then a 100-bin histogram formed for datapoints binned over every 2 seconds. Density plots were then formed by stacking these histograms vertically as shown in the main figures. For fitting the Ca^2+^ flux dynamics, the mean of the Indo-1 ratio at each time interval was first calculated and plotted. The light-induced decay of this signal was fit to the survival function of a gamma distribution over a range of shape parameters, which corresponds to the number of mass-action steps in a linear reaction scheme.

#### Phospho-ERK dynamics

To follow the dynamics of ERK phosphorylation, 0.5×10^6^ J.NFAT cells expressing the OptoCAR were conjugated with 1.5×10^6^ Raji-FRB^Ex^ cells as described above. Prior to the initiation of receptor activation, a sample was taken to define the 0-minute time-point. After this, the dimerizer was added at a final concentration of 2 μM and cells incubated at 37°C. After the required time, 100 μL of cell suspension was removed and immediately fixed in 500 μL 4% formaldehyde prewarmed to 37°C for 10 minutes. Samples were then kept on ice until timepoints had been acquired so that they could be further processed. Fixed samples were washed with 3 mL Flow wash buffer and centrifuged at 800×*g* for 3 minutes. The resuspended pellet was then vortexed while 500 μL Perm buffer III (cooled to −20°C) was added dropwise. After this, samples were left on ice for 30 minutes and then washed twice with 3 mL Flow wash buffer. The resuspended pellet was then incubated with the phospho-ERK1/2 (Thr^202^/Tyr^204^) antibody directly conjugated with Alexa Fluor 488 for 45 minutes at room temperature, with regular agitation. Stained cells were then washed with 3 mL Flow wash buffer and fixed with 500 μL Flow fix buffer before data acquisition by flow cytometry. Despite methanol fixation, the fluorescence from the iRFP713^+^ J.NFAT cells and mTagBFP^+^ Raji B cells was still readily detectable, allowing us to gate exclusively on cell conjugates for the subsequent analysis of phospho-ERK intensity.

#### Construction and calibration of the Optoplate illumination device

For a significant number of experiments in this work, we made use of an open-source illumination device, termed an ‘Optoplate’ that provides a way to independently illuminate all wells of a 96-well plate (Bugaj and Lim, 2019). We followed the authors’ protocol to build our own Optoplate using blue (470nm) LEDs in both positions under each well, with the following modifications. We increased the depth of the plate holder to 18mm and used 5mm thick Perspex squares as the diffusers in each position to improve the uniformity of illumination over the entire well. We also used a Raspberry Pi computer to communicate with the Arduino Micro onboard the Optoplate so that update commands for the intensity of each well could more easily be modulated. The Raspberry Pi used a simple CSV file written using MATLAB to send the intensity values for each well at an arbitrary refresh rate, normally every 10 seconds. This also allowed us to check on the status of the experiment by remote connection to the Optoplate whilst it was in the incubator.

Before any experiments were performed, the wells of the Optoplate were calibrated such that a defined pulse-width modulation (PWM) value applied to one well to control its intensity would be equivalent across the entire plate. We first imaged the Optoplate (without 96-well plate) using a gel-doc system at a range of PWM values, performed separately for the LED well pairs. We then used a MATLAB script to threshold the images to create well masks so that the mean intensity for each well could be calculated. We confirmed that all wells showed linearity of light intensity with PWM value before calculating calibration values so that the ‘dimmest’ well defined the maximum output and all other wells were proportionately scaled down. These calibration values were hardcoded onto the Arduino so that no calibration of the experimental program was required. We confirmed that this calibration process led to even illumination across the Optoplate, with a standard error of 0.4%.

#### FOS transcription factor dynamics

OptoCAR-T cells were conjugated as described above and dimerizer was added to 2 μM to initiate receptor signaling. For sample consistency, a stock of activated conjugates was prepared before adding 2 μM dimerizer and aliquoting into 24 wells of the Optoplate. We used Greiner μClear flat-bottomed plates for all experiments, which provided excellent light penetration and no spill over between wells. The first sample was then immediately taken to assay FOS levels at onset of activation. At the appropriate timepoint, all cells were removed from the required well (100 μL) and pipetted into an Eppendorf tube containing ice-cold 800 μL PBS. The tube was then centrifuged at 2000×*g* for 30 seconds, supernatant discarded and 100 μL RIPA buffer added to the pellet and gently mixed together. The lysed sample was incubated in ice for at least 30 minutes. When sample illumination was required, half of the Optoplate was switched on so that both ‘light’ and ‘dark’ samples could be collected simultaneously. Once all samples had been collected, the lysed cells were centrifuged at ~16,000×*g* for 10 minutes at 4°C. The supernatant (90 μL) was then pipetted into PCR tubes containing 42 μL of reduced LDS sample buffer and heated in a thermal cycler at 70°C for 10 minutes.

To detect total FOS and phosphorylated FOS protein levels, 15 μL of all samples was loaded onto two 26-well 8% Bis-Tris polyacrylamide gels and subjected to gel electrophoresis with a pre-stained protein ladder (BlueElf) in the end wells of the gel. Gels were then transferred to a nitrocellulose membrane using the iBlot2 dry blotting system. Blots were air-dried for 5 minutes before using No-Stain Protein Labelling Reagent to provide total protein normalization (TPN) loading control, following the manufacturer’s protocol. The blots were then immersed in blocking buffer (2.5% milk powder in Tris-buffered saline (TBS)) for 60 minutes. Primary antibody solutions were prepared in 10 mL blocking buffer supplemented with 0.1% Tween-20 (TBS-T/milk) using the FOS antibodies described in the Key Resources Table at manufacturer’s recommended dilution. Blots were incubated with antibody solutions overnight at 4°C on a rocking platform. After aspirating the primary antibody, blots were washed four times with TBS-T/milk before adding 10 mL secondary antibody solution containing CF790 anti-rabbit IgG(H+L) antibody at 1:10,000 dilution in TBS-T/milk. Blots were incubated for 60 minutes at room temperature on a rocking platform before being washed four times with TBS-T buffer. Blots were left in TBS and immediately imaged using the camera-based Azure 600 fluorescence detector.

To quantify the Western blot images, a line profile of each lane on the blots was measured using FIJI for both the TPN and FOS fluorescence channels and exported into MATLAB for further analysis. The band profiles were aligned through an automated script that used the TPN data to calculate the alignments that were applied to the FOS dataset. Background intensity was subtracted before numerical integration of the total peak intensities was calculated. For the TPN data, almost all the lane intensity was included in this integration to give the most reliable measure of protein loading. To calculate the ‘active’ fraction of FOS when required, a binary gate was applied to the histogram of the lane profile to define the higher molecular weight part of profile. This dataset was normalized between the 0- and 15-minute timepoints.

#### Measurement of downstream OptoCAR output

To provide more robust control over downstream signaling, we used the V416L variant of LOV2 (Wang et al., 2016) in the OptoCAR, which has a slower dark reversion rate and requires lower light intensity to maintain signaling quiescence. This had the benefit of keeping the cell conjugates healthier over the 24-hour illumination period by removing any phototoxicity effect. OptoCAR-T cells were conjugated with ligand-presenting cells as described above. For reproducibility, a large stock of cell conjugates was prepared using 8×10^6^ OptoCAR-T cells and 15×10^6^ Raji-FRB^Ex^ cells, before adding 2 μM dimerizer and aliquoting 200 μL of conjugates into all wells of a 96-well plate, which corresponded to 8×10^4^ OptoCAR-T cells and 1.5×10^5^ Raji-FRB^Ex^ cells per well. We used Greiner μClear flat-bottomed plates for all experiments, which provided excellent light penetration and no spill over between wells. The Optoplate was then placed in a 37°C/5% CO_2_ incubator and run for 24 hours with individual well illumination profiles appropriately programmed depending on the required experiment. Signaling (‘dark’) pulses were 15, 30, 60, 72, 90 or 120 minutes in duration and the inter-pulse times when needed were calculated between a duty cycle of 100% and 25%. Duplicates wells of constant dark, constant light and 6 hours continuous dark were included for all experiments. The illumination profiles for each experiment were randomized across the Optoplate to remove any potential positional effect that might have biased the results. On completion of the experiment, samples were stained for CD69 surface expression in 96 well round bottom plates and analyzed by flow cytometry as described above.

Experimental datasets were analyzed using FlowJo to gate on OptoCAR-T cells and quantify the geometric mean of the GFP and CD69 (Alexa Fluor 647) expression distributions. These values were then normalized between 6 hours of continuous signaling and the constant light control and plotted against the inter-pulse interval.

#### Reverse-transcription quantitative PCR (RT-qPCR) to measure mRNA levels

OptoCAR-T cells and Raji cells were conjugated similar to other assays such that each well contained 3.2×10^5^ OptoCAR-T cells conjugated with 5.6×10^5^ Raji-FRB^Ex^. A pooled stock of activated conjugates was prepared before adding 2 μM dimerizer to initiate the receptor-signaling and then aliquoting cell conjugates into 26 wells of Greiner μClear flat-bottomed 96 well plate. Eleven wells were illuminated after 3 hours, with the remainder of wells kept in the dark for the entire assay. A volume of 100 μL was removed for each sample and put into 1.5ml Eppendorf tube containing 800 μL PBS. The tube was centrifuged at 500×*g* for 30 seconds, the supernatant was discarded before placing tube in a pre-cooled metal block in dry ice to rapidly freeze the cell pellets. Other samples were taken out in same manner at specified time points.

Frozen cell pellets were thawed on ice, then total RNA was extracted using RNeasy Plus Mini Kit (Qiagen) following manufacturer’s protocol, with Qiashredder columns used to homogenize samples and reduce viscosity. The total RNA concentration was measured using NanoDrop Spectrophotometer (GeneFlow) at 260nm and RNA quality was assessed by measuring 260 / 280 nm absorbance ratio. cDNA was reverse transcribed from 2 μg of RNA using SuperScript IV VILO Master Mix with ezDNase Enzyme (Invitrogen) following manufacturer’s protocol; a ‘No-RT’ control sample was always prepared in conjunction.

The relative mRNA levels of defined genes were analyzed by quantitative real-time PCR, with *GAPDH* and *PGK1* used as reference housekeeping genes. The 20 μL PCR reaction mix was prepared with 10 μL TaqMan Advanced Fast Master Mix, 1 μL TaqMan Assay for each gene, 7 μL nuclease free water and 2 μL cDNA sample. The reaction mix were pipetted into 96 well PCR plates and sealed with optical Cap strips. The plate was vortexed to mix the reagents and then briefly centrifuged to bring the reaction mix to the bottom of each tube and eliminate air bubbles. The plate was then loaded onto an Agilent Mx3005P Real-Time PCR System and run using cycling conditions as follows: UNG(uracil-N-glycosylase) incubation at 50°C for 2 minutes, polymerase activation at 95°C for 2 minutes followed by 40 PCR cycles of denaturation at 95°C for 3 seconds and annealing/extension at 60°C for 30 seconds.

The cycle threshold (Ct) value for each PCR reaction was extracted from the fluorescent traces and converted to a relative quantity (RQ) by taking the difference between each timepoint and the initial value at the start of the experiment. We confirmed that the efficiencies of each TaqMan assay was ~100% over the range of extracted Ct values. The RQ values were then corrected (cRQ) for total input using the geometric mean of the paired RQ values for *GAPDH* and *PGK1* housekeeping genes. These values were then normalized over [0,1] using the mean cRQ values between 180 and 240 minutes for each dataset and this is what is plotted in the main figure. All experiments are represented as the mean of three biological replicates using single or double technical replicates.

#### Cellular response to varying signaling potency

To titrate the signaling ‘strength’ within OptoCAR-T cells, the Optoplate was used to continuously illuminate cell conjugates over a range of intensities, with the experiment set up in an equivalent manner to the previous sections. The gated sample data from FlowJo was then imported into MATLAB for further analysis. The distributions of fluorescence intensity from both NFAT-mediated GFP expression and CD69 upregulation were constructed for each well before combining vertically to build a density plot with downstream output expression against illumination intensity.

#### Microscopy of OptoCAR-T cell conjugate activation

To quantify expression of NFAT-mediated GFP expression over time from individual cell conjugates, we used live-cell spinning disk confocal microscopy. OptoCAR-T cell conjugates were prepared as described above but using the LOV2 C450G mutation so that the OptoCAR remained in the signaling-competent state even during imaging at wavelengths < 500nm that would normally disrupt it. Conjugated cells were pipetted into a fibronectin-coated imaging dish and placed on the microscope maintained at 37°C and 5% CO2. A 100× oil-immersed objective was used to image the J.NFAT cells (iRFP713: 640nm Ex; 708±37nm Em), Raji-FRBEx cells (mTagBFP: 405 Ex; 460±30nm Em), OptoCAR construct (mScarlet: 561nm Ex; 607±18nm Em) and GFP (488 Ex; 525±25nm Em) every 10 minutes for 24 hours. In order to image a significant number of conjugates at this high resolution, we imaged a 12×12 grid of sample regions (512×512 px/region) with 10% overlap and then stitched the separate stacks together using ‘Grid Stitching’ in FIJI. This stitched image stack was then split into 9 sub-stacks to facilitate analysis and five of these regions maintained enough conjugates over 24 hours to quantify the mean GFP intensity over time.

#### Measurement of OptoCAR^CD19^ activation

Human primary CD4+ T cells were transduced with the anti-CD19 OptoCAR (OptoCAR^CD19^) construct as previously described (James, 2018). The OptoCAR^CD19^-T cells were conjugated with Raji B cells as for the OptoCAR experiments above (but without the dimerizer addition) and various illumination profiles were patterned onto the cells over 24 hours using the OptoPlate. On completion of the experiment, samples were stained for CD3 and CD20 to differentiate the two celltypes by flow cytometry, as well as CD137 to quantify T cell activation. All OptoCAR^CD19^-positive T cells (as gated by mScarlet fluorescence) were found to remain conjugated with B cells so this population was quantified for CD137 expression. These values were then normalized between 6 hours of continuous signaling and the constant light control and plotted against the inter-pulse interval.

### QUANTIFICATION AND STATISTICAL ANALYSIS

The quantification of datasets is described in the methods sections above for clarity. Where mean values are reported in the main text, the standard error of measurement is also reported, along with the number of biological replicates used to calculate this value. For some of the decay rates presented in the main text, a 95% confidence interval is also provided, using the t-distribution to appropriately bound the interval.

## SUPPLEMENTAL INFORMATION LEGENDS

**Figure S1. Quantifying the intracellular Ca^2+^ flux in conjugated OptoCAR-T cells. Related to Figure 2.**

(A-C) Flow cytometry data of Jurkat T cells expressing the OptoCAR construct after lentiviral transduction. The distribution of mScarlet fluorophore genetically fused to OptoCAR is shown (A), along with surface staining for the receptor using an anti-CD86 antibody (B). The bivariate plot of these two distributions shows that they are extremely well correlated (C), demonstrating mScarlet intensity is a good marker for OptoCAR cell-surface expression.

(D, E) Conjugates formed between Jurkat (J.NFAT; IRFP713^+^) and Raji FRB^Ex^ cells (mTagBFP^+^) can be directly gated on during acquisition of flow cytometry data. Simply mixing the two cell-types together produces essentially no double-positive events (D) but allowing them to interact at high density at 37°C for 10 minutes creates a distinct and readily observable population of cell conjugates (E).

(F) Photo of the custom-built illumination device that can expose the sample to blue light whilst maintaining it at 37°C on the flow cytometer.

(G, H) Repeating the Ca^2+^ flux assay described in the main text with a variant of the LOV2 domain in the OptoCAR that is unresponsive to blue-light illumination (OptoCAR^C450G^) drives increased intracellular [Ca^2+^] independently from the illumination state of the sample (G). Conversely, the complementary mutation of the LOV2 domain that maintains it in the ‘light’ state (OptoCAR^I539E^) cannot drive signaling even in the dark (H).

(I) The dataset in Figure 3L was fitted using the survival function of a gamma distribution with the shape parameter varying, which is analogous to the number of reaction steps. By minimizing the sum of the squared residuals (SSR) for each fit, we found that the dataset was fitted best to a model with five steps.

**Figure S2. Uncropped Western blots of FOS datasets and conjugation time course. Related to Figure 4.**

(A) Uncropped fluorescent Western blots from data presented in Figure 5A. The accompanying total protein normalization blot for this blot is also provided in the lower image. Molecular weight markers were determined from an additional fluorescence channel.

(B) Uncropped fluorescent Western blots from data presented in Figure 5B. The accompanying total protein normalization blot for this blot is also provided in the lower image. Molecular weight markers were determined from an additional fluorescence channel.

(C) The fraction of Jurkat T cells conjugated with Raji B cells over time was measured by flow cytometry. Dimerizer was added to the cells at the start of experiment, which drives more stable conjugates. Each line represents a replicate test.

**Figure S3. Downstream output from OptoCAR-T cells can be modulated by illumination. Related to Figure 5.**

(A) The ability of illumination to modulate NFAT-mediated GFP expression from OptoCAR-T cells was tested by either activating cells continuously in the dark or light state (denoted in boxes above plots). Vehicle control (no dimerizer) was used to assess baseline expression. Plots show flow cytometry data with GFP intensity against OptoCAR expression.

(B) The ability of illumination to modulate CD69 upregulation from OptoCAR-T cells was tested by either activating cells continuously in the dark or light state (denoted in boxes above plots). Vehicle control (no dimerizer) was used to assess baseline expression. Plots show flow cytometry data with surface CD69 intensity against OptoCAR expression.

(C) The complete dataset of the plot shown in Figure 5B. Plot shows the geometric mean of GFP expression over a 24-hour range of signal pulses, normalized to output at 24 hours. Error bars show mean ± SEM (n=3).

(D) The complete dataset of the plot shown in Figure 5D. Plot shows the geometric mean of CD69 upregulation over a 24-hour range of signal pulses, normalized to output at 24 hours. Error bars show mean ± SEM (n=3).

(E) Flow cytometry plots showing the distribution of NFAT-mediated GFP expression after different single pulse lengths (denoted in boxes above plots). A subset of pulse lengths is shown, ranging from 15 minutes to 24 hours. These distributions were used to build the density plot in Figure 5A.

(F) Flow cytometry plots showing the distribution of CD69 upregulation after different single pulse lengths (denoted in boxes above plots). A subset of pulse lengths is shown, ranging from 15 minutes to 24 hours. These distributions were used to build the density plot in Figure 5C.

(G) Combined plot of all datasets from Figure 5F but only over inter-pulse interval times up to 20 minutes, with left panel showing the unaltered data and the right panel showing the datasets normalized to minimal observed output for each dataset. The plots for 90 and 120 minute pulse duration have been omitted as there was insufficient dynamic range in these datasets.

(H) Combined plot of all datasets from Figure 5G but only over inter-pulse interval times up to 20 minutes, presented equivalently to (G).

**Figure S4. Graded illumination of OptoCAR-T cells drives both analogue and digital output. Related to Figure 5.**

(A) Plot showing the linearity between the pulse width modulation (PWM) value that controlled LED intensity and the corresponding light output on the OptoPlate, which was used to calibrate the OptoPlate in subsequent experiments.

(B) Density plot of NFAT-mediated GFP expression as a function of light intensity continuously applied to OptoCAR-T cell conjugates over 24 hours. Plot is composed from 85 individual histograms.

(C) Density plot of CD69 upregulation as a function of light intensity continuously applied to OptoCAR-expressing T cells, from same dataset as in (A).

(D) Density plot of NFAT-mediated GFP expression as a function of light intensity continuously applied to OptoCAR-expressing T cells. The new range of illumination was chosen to better resolve bistable region in (B). Plot is composed from 96 individual histograms.

**Figure S5. Vector construction and protein sequence of OptoCARs. Related to STAR Methods.**

(A) Schematic showing the various components part of the OptoCAR construct as they are arranged in the lentiviral vector used to express the receptor in T cells. Relevant restriction sites are highlighted. Myr denotes myristoylation sequence and SP is signal peptide sequence.

(B) Protein sequence of OptoCAR, using colors from (A) to denote different regions. Point mutations in LOV2 domain discussed in STAR methods are highlighted red.

(C) Oligonucleotide sequences used in construction of OptoCAR vectors.

(D) Schematic showing the various components part of the OptoCAR^CD19^ construct as they are arranged in the lentiviral vector used to express the receptor in T cells. Relevant restriction sites are highlighted. Myr denotes myristoylation sequence.

(E) Protein sequence of OptoCAR^CD19^, using colors from (D) to denote different regions.

**Movie S1. Dynamics of light-induced dissociation of OptoCAR. Related to Figure 2.**

Movie showing the membrane-bound form of the OptoCAR expressed in HEK293T cells being reversibly dissociated from extracellular domain by single-cell illumination (white box). The intracellular part of the OptoCAR used in this experiment was not myristoylated so it localized to the cytoplasm on dissociation, to aid visualization of OptoCAR dissociation.

**Movie S2. NFAT-mediated GFP expression in a single cell conjugate. Related to Figure 6.**

Movie following a single J.NFAT cell expressing the OptoCAR (red channel) initiating contact with a Raji-FRB^Ex^ cell (blue channel) and becoming activated, quantified by the increased GFP fluorescence (green channel) mediated by NFAT transcriptional activity. During the movie, the T cell engages with an alternative Raji-FRB^Ex^ cell but never becomes dissociated at any point. The C450G variant of the OptoCAR was used, which is unresponsive to light modulation, so that GFP imaging did not disrupt receptor signaling. Timestamps show hours elapsed.

**Movie S3. Effect of graded illumination of OptoCAR-T cell conjugates on downstream output. Related to Figure 7.**

A sequence of histograms of either NFAT-mediated GFP expression (left panel) or CD69 downregulation (right panel) from OptoCAR-T cell conjugates under continuous illumination at varying intensity (blue scale bar). These distributions were used to build the density plots in Figures 7A and 7B.

**Movie S4. Effect of high-resolution graded illumination of OptoCAR-T cell conjugates on NFAT-mediated GFP expression. Related to Figure 7.**

A sequence of histograms of NFAT-mediated GFP expression from OptoCAR-T cell conjugates under continuous illumination at varying intensity (blue scale bar). These distributions were used to build the density plot in Figure 7C.

