## Supplementary Figures (Combined) for "Quantifying signal persistence in the T cell signaling network using an optically controllable antigen receptor"

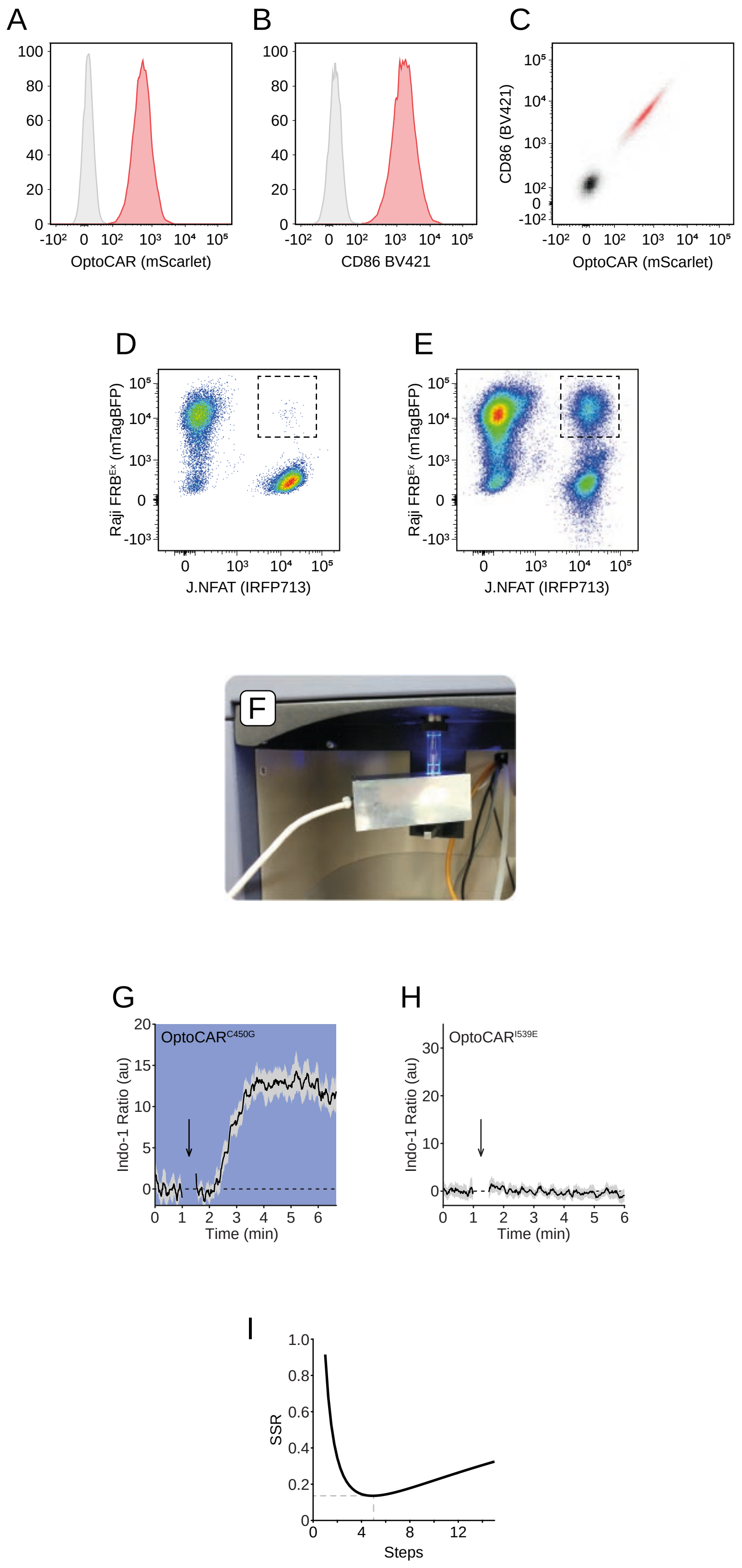
A
B
C
100
80
60
40
20
0
-10²
0
10²
10³
10⁴
10⁵
OptoCAR (mScarlet)
CD86 BV421
CD86 (BV421)
D
E
Raji FRBᴱˣ (mTagBFP)
-10³
J.NFAT (IRFP713)
F
G
H
OptoCAR^C450G
OptoCAR^I539E
Indo-1 Ratio (au)
Time (min)
I
SSR
Steps

Figure S1

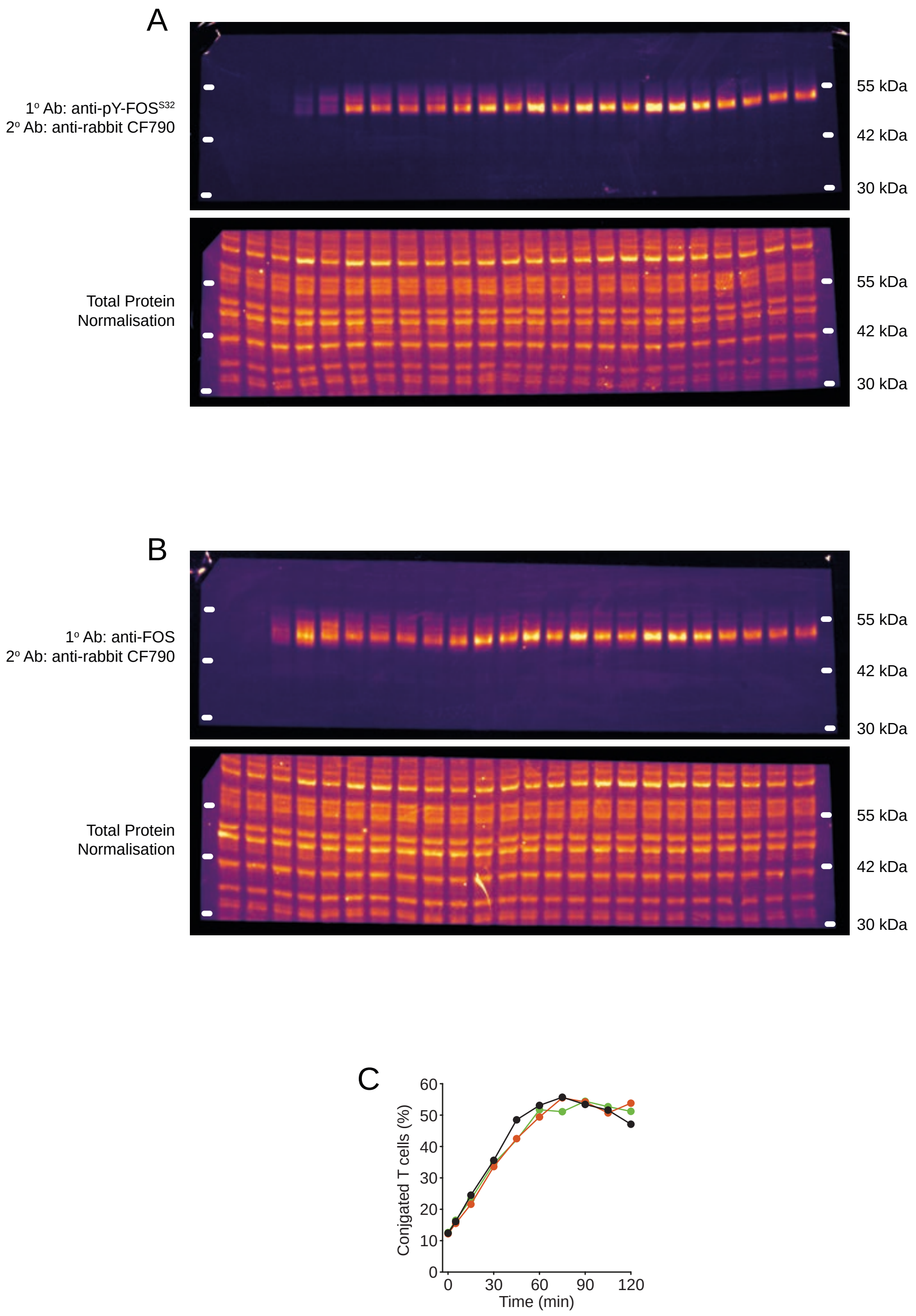
A
1º Ab: anti-pY-FOS$^{S32}$
2º Ab: anti-rabbit CF790
55 kDa
42 kDa
30 kDa
Total Protein
Normalisation
55 kDa
42 kDa
30 kDa
B
1º Ab: anti-FOS
2º Ab: anti-rabbit CF790
55 kDa
42 kDa
30 kDa
Total Protein
Normalisation
55 kDa
42 kDa
30 kDa
C
Conjugated T cells (%)
0
10
20
30
40
50
60
0
30
60
90
120
Time (min)

Figure S2

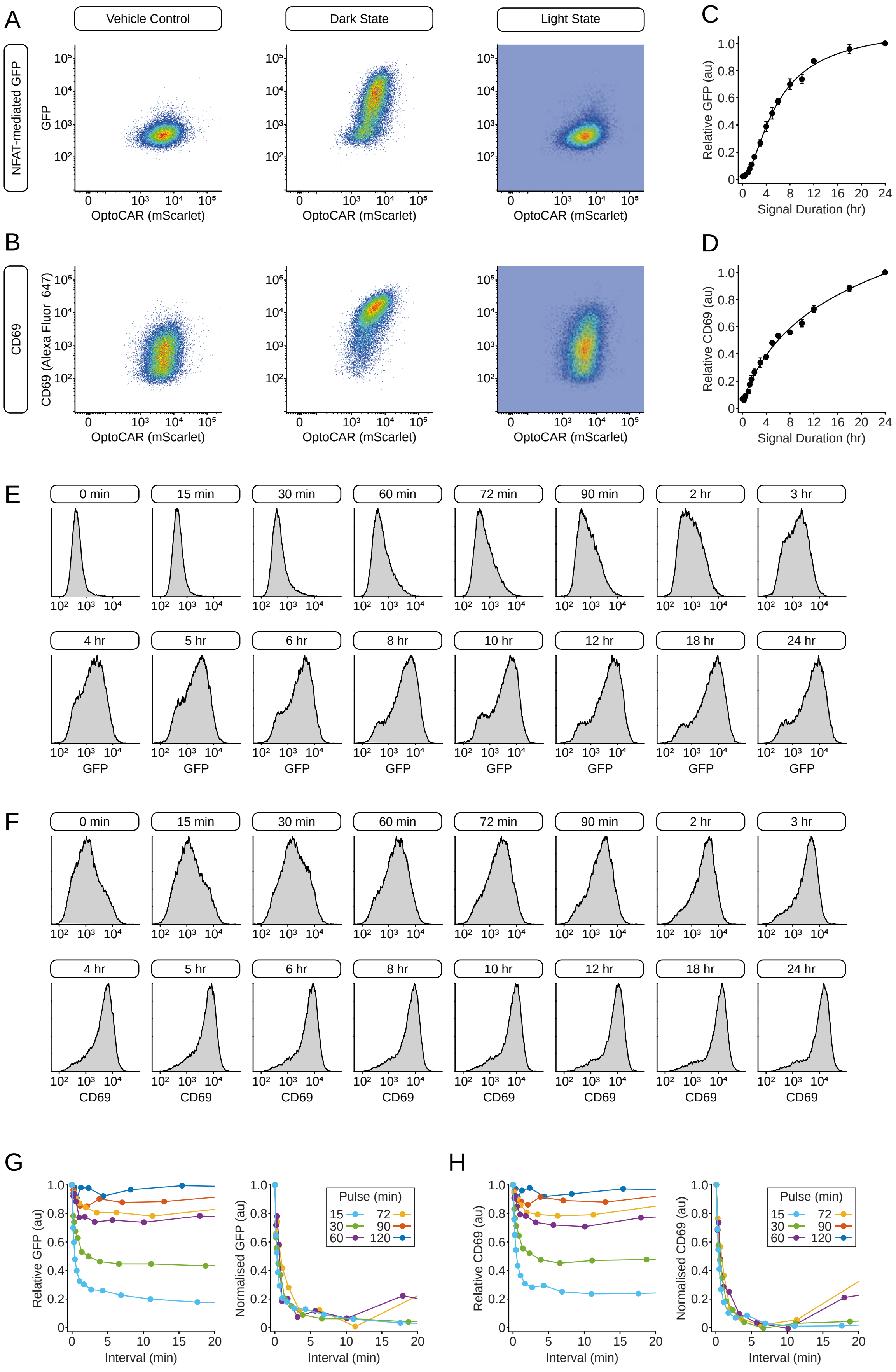

A
Vehicle Control
Dark State
Light State
NFAT-mediated GFP
GFP
OptoCAR (mScarlet)
C
Relative GFP (au)
Signal Duration (hr)
B
CD69
CD69 (Alexa Fluor 647)
D
Relative CD69 (au)
E
0 min
15 min
30 min
60 min
72 min
90 min
2 hr
3 hr
4 hr
5 hr
6 hr
8 hr
10 hr
12 hr
18 hr
24 hr
F
G
Relative GFP (au)
Normalised GFP (au)
Interval (min)
Pulse (min)
H
Relative CD69 (au)
Normalised CD69 (au)


### Figure S3

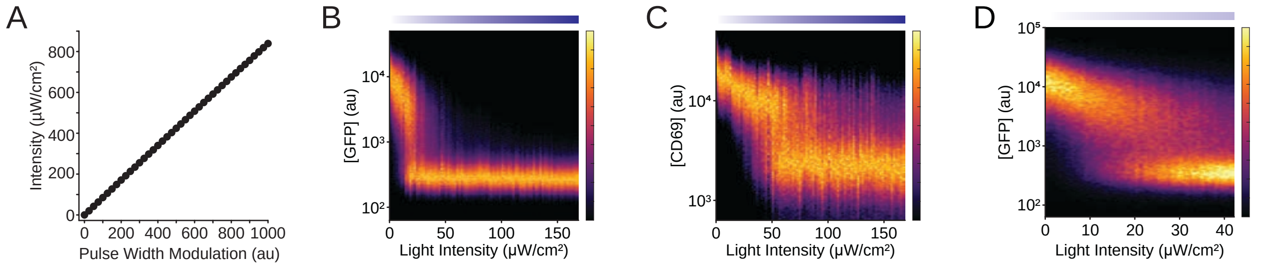
A
B
C
D
Intensity (µW/cm²)
800
600
400
200
0
0 200 400 600 800 1000
Pulse Width Modulation (au)
[GFP] (au)
10⁴
10³
10²
0 50 100 150
Light Intensity (µW/cm²)
[CD69] (au)
10⁴
10³
0 50 100 150
Light Intensity (µW/cm²)
[GFP] (au)
10⁵
10⁴
10³
10²
0 10 20 30 40
Light Intensity (µW/cm²)

Figure S4

A

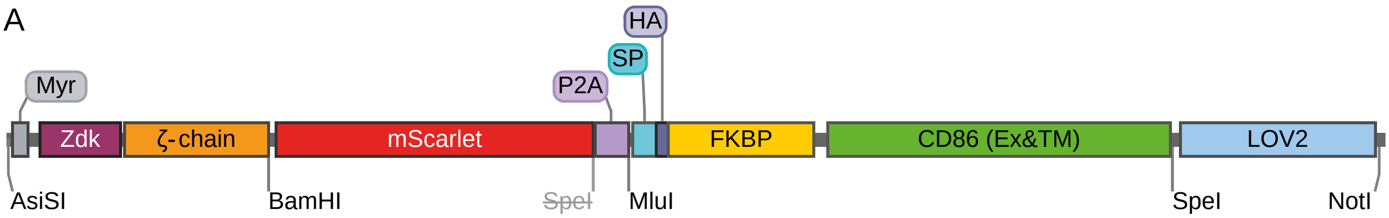


B

```
  1 MGCGCSSHPE DDGGSGGSGG SMVDNKFNKE KTRAGAEIHS LPNLNVEQKF AFIVSLFDDP SQSANLLAEA  70
 71 KKLNDAQAPK TSADAPAYQQ GQNQLYNELN LGRREEYDVL DKRRGRDPEM GGKPQRRKNP QEGLYNELQK 140
141 DKMAEAYSEI GMKGERRRGK GHDGLYQGLS TATKDTYDAL HMQALPPRDP PVATMVSKGE AVIKEFMRFK 210
211 VHMEGSMNGH EFEIEGEGEG RPYEGTQTAK LKVTKGGPLP FSWDILSPQF MYGSRAFTKH PADIPDYYKQ 280
281 SFPEGFKWER VMNFEDGGAV TVTQDTSLED GTLIYKVKLR GTNFPPDGPV MQKKTMGWEA STERLYPEDG 350
351 VLKGDIKMAL RLKDGGRYLA DFKTTYKAKK PVQMPGAYNV DRKLDITSHN EDYTVVEQYE RSEGRHSTGG 420
421 MDELYKTSYQ GPGATNFSLL KQAGDVEENP GPTRMGVKVL FALICIAVAE AKYPYDVPDY AGVQVETISP 490
491 GDGRTFPKRG QTCVVHYTGM LEDGKKVDSS RDRNKPFKFM LGKQEVIRGW EEGVAQMSVG QRAKLTISPD 560
561 YAYGATGHPG IIPPHATLVF DVELLKLEGG SGGSGGSLKI QAYFNETADL PCQFANSQNQ SLSELVVFWQ 630
631 DQENLVLNEV YLGKEKFDSV HSKYMGRTSF DSDSWTLRLH NLQIKDKGLY QCIIHHKKPT GMIRIHQMNS 700
701 ELSVLANFSQ PEIVPISNIT ENVYINLTCS SIHGYPEPKK MSVLLRTKNS TIEYDGIMQK SQDNVTELYD 770
771 VSISLSVSFP DVTSNMTIFC ILETDKTRLL SSPFSIELED PQPPPDHIPW ITAVLPTVII CVMVFCLILW 840
841 KWKKKKRPRT SGGSGSLATT LERIEKNFVI TDPRLPDNPI IFASDSFLQL TEYSREEILG RNCRFLQGPE 910
911 TDRATVRKIR DAIDNQTEVT VQLINYTKSG KKFWNLFHLQ PMRDQKGDVQ YFIGVQLDGT EHVRDAAERE 980
981 GVMLIKKTAE NIDEAAKEL*                                                       999
```

C

```
pr1: 5' gtagactagtggcggaagcggaagtttggctactacacttgaacgtattg 3'
pr2: 5' ctactagcggccgctcaaagttcttttgccgcctcatc 3'
pr3: 5' tagtaggcgatcgccaccatgggctgtggctgcagc 3'
pr4: 5' cgccagggccctggtaactGgtcttgtacagctcgtccatgc 3'
pr5: 5' gcatggacgagctgtacaagacCagttaccagggccctggcg 3'
pr6: 5' tagtaggcgatcgccacctggtggataacaaattcaataaagaaaag 3'
pr7: 5' ctactaggatcccgagggggcagggcc 3'
```

D

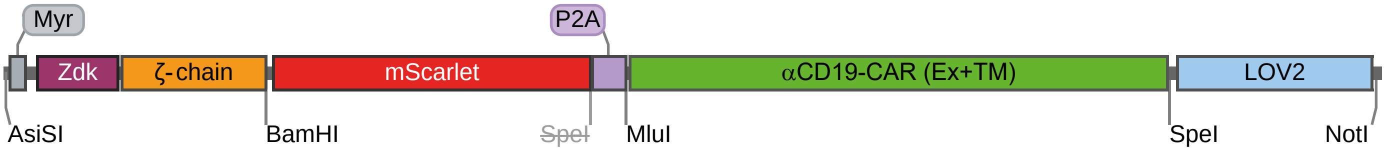


E

```
  1 MGCGCSSHPE DDGGSGGSGG SMVDNKFNKE KTRAGAEIHS LPNLNVEQKF AFIVSLFDDP SQSANLLAEA  70
 71 KKLNDAQAPK TSADAPAYQQ GQNQLYNELN LGRREEYDVL DKRRGRDPEM GGKPQRRKNP QEGLYNELQK 140
141 DKMAEAYSEI GMKGERRRGK GHDGLYQGLS TATKDTYDAL HMQALPPRDP PVATMVSKGE AVIKEFMRFK 210
211 VHMEGSMNGH EFEIEGEGEG RPYEGTQTAK LKVTKGGPLP FSWDILSPQF MYGSRAFTKH PADIPDYYKQ 280
281 SFPEGFKWER VMNFEDGGAV TVTQDTSLED GTLIYKVKLR GTNFPPDGPV MQKKTMGWEA STERLYPEDG 350
351 VLKGDIKMAL RLKDGGRYLA DFKTTYKAKK PVQMPGAYNV DRKLDITSHN EDYTVVEQYE RSEGRHSTGG 420
421 MDELYKTSYQ GPGATNFSLL KQAGDVEENP GPTRMALPVT ALLLPLALLL HAARPDIQMT QTTSSLSASL 490
491 GDRVTISCRA SQDISKYLNW YQQKPDGTVK LLIYHTSRLH SGVPSRFSGS GSGTDYSLTI SNLEQEDIAT 560
561 YFCQQGNTLP YTFGGGTKLE ITGGGGSGGG GSGGGGSEVK LQESGPGLVA PSQSLSVTCT VSGVSLPDYG 630
631 VSWIRQPPRK GLEWLGVIWG SETTYYNSAL KSRLTIIKDN SKSQVFLKMN SLQTDDTAIY YCAKHYYYGG 700
701 SYAMDYWGQG TSVTVSSTTT PAPRPPTPAP TIASQPLSLR PEACRPAAGG AVHTRGLDFA CDIYIWAPLA 770
771 GTCGVLLLSL VITLYCKKKK RPRTSGGSGS LATTLERIEK NFLITDPRLP DNPIIFASDS FLQLTEYSRE 840
841 EILGRNCRFL QGPETDRATV RKIRDAIDNQ TEVTVQLINY TKSGKKFWNL FHLQPMRDQK GDVQYFIGVQ 910
911 LDGTEHVRDA AEREGVMLIK KTAENIDEAA KEL*                                       943
```

Figure S5
