## Supplementary figures and images for "Quantifying signal persistence in the T cell signaling network using an optically controllable antigen receptor"

### Supplementary Movie 2

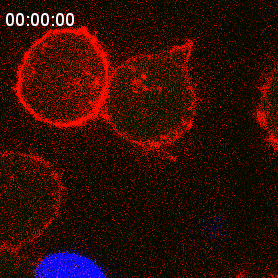

### Supplementary Movie 3

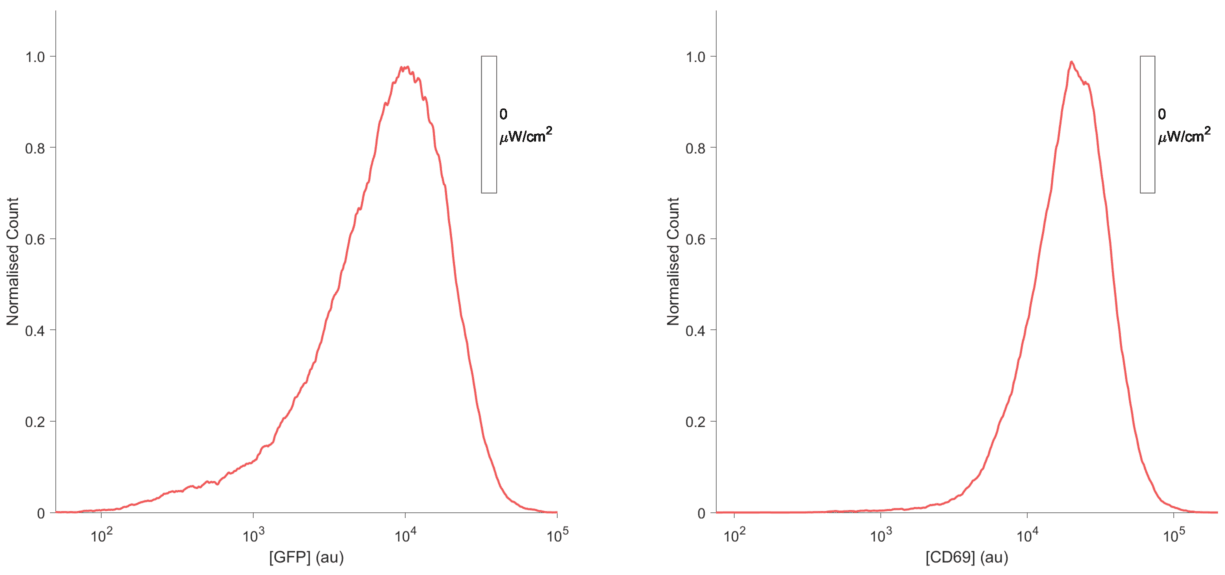

### Supplementary Movie 4

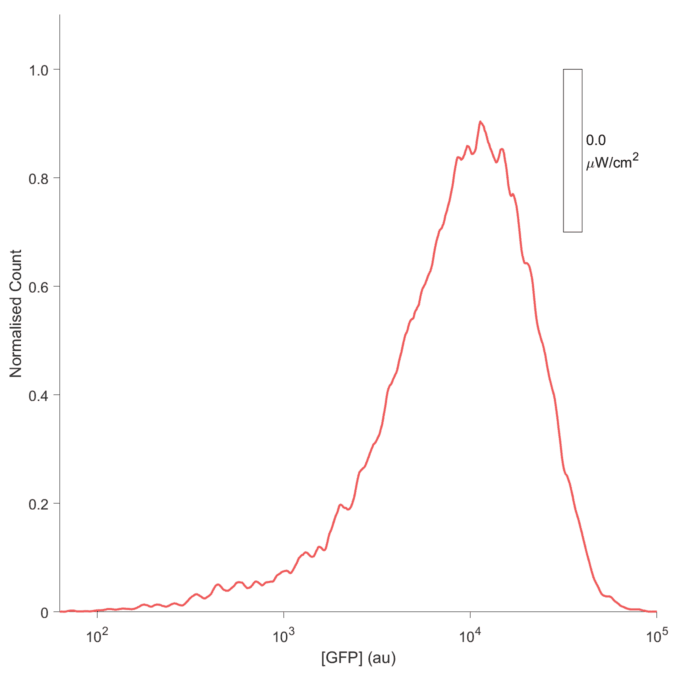
